# *Pseudomonas aeruginosa* FpvB is a high-affinity transporter for xenosiderophores ferrichrome and ferrioxamine B

**DOI:** 10.1101/2022.09.20.508722

**Authors:** Derek C. K. Chan, Lori L. Burrows

## Abstract

Iron is essential for many biological functions in bacteria but its poor solubility is a limiting factor for growth. Bacteria produce siderophores, soluble natural products that bind iron with high affinity, to overcome this challenge. Siderophore-iron complexes return to the cell through specific outer-membrane transporters. The opportunistic pathogen *Pseudomonas aeruginosa* makes multiple transporters that recognize its own siderophores, pyoverdine and pyochelin, and xenosiderophores produced by other bacteria, which gives it a competitive advantage. Some antibiotics exploit these transporters to bypass the membrane to reach their intercellular targets – including the thiopeptide antibiotic, thiostrepton (TS), which uses the pyoverdine transporters FpvA and FpvB to cross the outer membrane. Here, we assessed TS susceptibility in the presence of various siderophores and discovered that ferrichrome and ferrioxamine B antagonized TS uptake via FpvB. Unexpectedly, we found that FpvB transports ferrichrome and ferrioxamine B with higher affinity than pyoverdine. Site-directed mutagenesis of FpvB coupled with competitive growth inhibition and affinity label quenching studies suggested that the siderophores and antibiotic share a common binding site in an aromatic pocket formed by the plug and barrel domains, but have differences in their binding mechanism and molecular determinants for uptake. This work describes an alternative uptake pathway for ferrichrome and ferrioxamine B in *P. aeruginosa* and emphasizes the promiscuity of siderophore transporters, with implications for Gram-negative antibiotic development via the Trojan horse approach.

## INTRODUCTION

Iron is an essential micronutrient for bacteria, but has poor aqueous solubility at neutral pH and consequently, low bioavailability (Klebba et al., 2021; Stefánsson, 2007). At sites of infection, bacteria also compete with host-defense proteins that sequester iron. To overcome these limitations, Gram-negative bacteria secrete siderophores, small molecules with high affinity for iron. Once outside the cell, siderophores scavenge iron and return through specific outer-membrane transporters on the cell surface (Noinaj et al., 2010). The architecture of siderophore transporters is conserved, consisting of a 22-stranded beta-barrel with a plug domain that occludes the lumen to prevent passive diffusion (Noinaj et al., 2010). The extracellular loops of the transporters are important for siderophore recognition and uptake. The periplasmic N-terminus contains a short motif known as the TonB box (Josts et al., 2019; Moynié et al., 2019; Noinaj et al., 2010), which interacts with the inner membrane protein TonB. Together with the inner membrane proteins ExbB-ExbD, TonB harnesses the proton motive force to actively transport ligands through the transporters via a mechanism that remains incompletely understood (Klebba et al., 2021; Noinaj et al., 2010). Although TonB-dependent transporters (TBDTs) are considered ligand-specific, they can be exploited by antimicrobial compounds, bacteriophages, and bacteriocins for uptake, making them of interest for drug delivery across the outer membrane of Gram negatives (Chan and Burrows, 2021; Ferguson et al., 2000; Luna et al., 2020; Luscher et al., 2018; Mathavan et al., 2014; Rabsch et al., 2007; Ranieri et al., 2019; Salomon and Farias, 1993; White et al., 2017).

The opportunistic bacterial pathogen, *Pseudomonas aeruginosa* encodes ~35 TBDTs for different ligands including siderophores, cobalamin, and other metal complexes (Hancock and Brinkman, 2003; Klebba et al., 2021; Llamas et al., 2006; Luscher et al., 2018; Moynié et al., 2019; Ochsner et al., 2000). *P. aeruginosa* makes two main siderophores, pyoverdine and pyochelin, which are taken up via FpvA and FpvB, and FptA, respectively (Cobessi et al., 2005; Ghysels et al., 2004; Heinrichs et al., 1991; Meyer et al., 2002; Sokol, 1987) (Figure 1). Pyoverdine has higher affinity for iron than pyochelin (Albrecht-Gary et al., 1994; Braud et al., 2009) and has roles in tolerance to antibiotics, biofilm formation, and virulence factor production (Banin et al., 2005; Lamont et al., 2002). *P. aeruginosa* can also take up siderophores produced by other microorganisms, including ferrioxamine E and B (produced by *Streptomyces spp.*) and ferrichrome (produced by fungal species) via the FoxA and FiuA TBDTs, respectively (Klebba et al., 2021; Llamas et al., 2006; Normant et al., 2020) (Figure 1).

**Figure 1.**
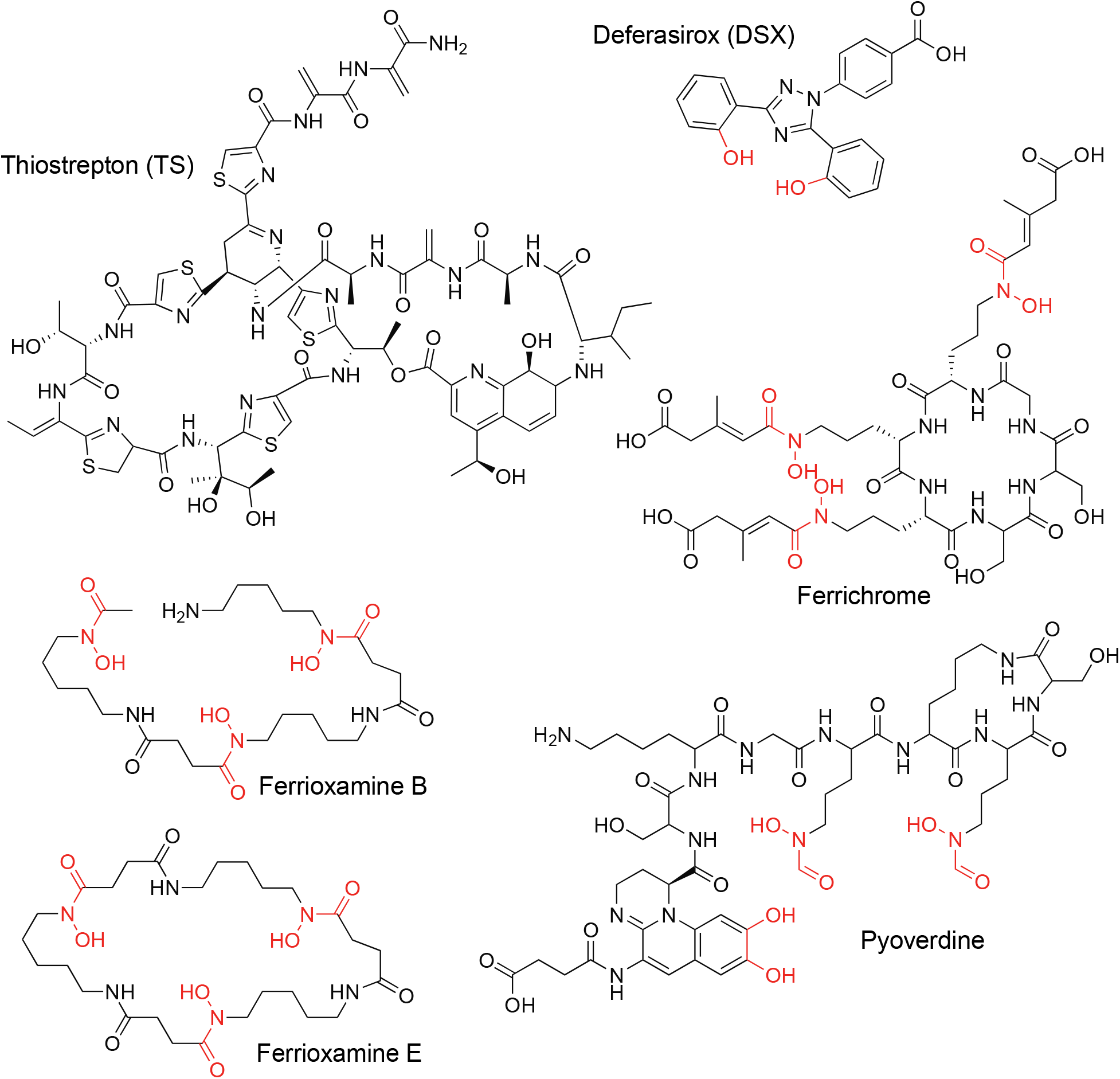
Structures of compounds used in this study. Iron chelating groups are highlighted in red.

Understanding the range of ligands that can be taken up by TBDTs is important as there is growing interest in designing novel antibiotics that can exploit these transporters for uptake. However, our understanding of the substrate range for individual TBDTs is lacking. Even for those with known ligands, there may be substrates that have yet to be discovered. For example, ferrioxamine E exclusively uses FoxA to enter *P. aeruginosa* (Normant et al., 2020). However, after *foxA* or *fiuA* are deleted, the bacteria still grow in iron-limited media when supplied with ferrioxamine B or ferrichrome, respectively, suggesting these siderophores can also be recognized by other transporters (Normant et al., 2020).

Previously we showed that the large thiopeptide antibiotics thiostrepton (TS) and thiocillin (TC) use TBDTs to enter *P. aeruginosa* to access their cytoplasmic target, the ribosome. TS exploits the pyoverdine transporters, FpvA and FpvB, while thiocillin uses the ferrioxamine transporter, FoxA (Figure 1) (Chan et al., 2020; Chan and Burrows, 2021; Ranieri et al., 2019). Here we further characterized the interaction of TS with the pyoverdine transporters. We discovered that the secondary transporter FpvB has high affinity for ferrichrome and ferrioxamine B but is a poor pyoverdine transporter. FpvB is a promiscuous transporter as it can recognize structurally distinct ligands using different binding modes. Overall, this work expands our understanding of TBDT function and fills in the missing details of ferrichrome and ferrioxamine B uptake in *P. aeruginosa*.

## RESULTS

### Exogenous pyoverdine poorly rescues iron-restricted growth of a PA14 Δ*fpvA* mutant

Previously we showed that TS synergized with the FDA-approved iron chelator deferasirox (DSX) against PA14, and that susceptibility required the FpvA and FpvB pyoverdine transporters (Figure 1) (Ranieri et al., 2019). DSX lacks antimicrobial activity against wild type (WT) cells since they produce pyoverdine, which competes with DSX for iron. Therefore, WT PA14, a Δ*fpvA* mutant, and a Δ*fpvB* mutant are each expected to be susceptible to TS and grow in the presence of DSX since they still encode functional pyoverdine transporters. A Δ*fpvA* Δ*fpvB* mutant is resistant to TS and inhibited by DSX (Chan and Burrows, 2021; Ranieri et al., 2019).

We first confirmed these phenotypes via minimal inhibitory concentration (MIC) assays. The WT and Δ*fpvA* mutant were susceptible to TS in iron-limited medium (Figure 2A). The Δ*fpvB* mutant was resistant to TS although its growth was reduced at the maximum soluble TS concentration of 17 μg/mL. The Δ*fpvA* Δ*fpvB* mutant was resistant to TS with no observable decrease in growth. The MIC of DSX against the Δ*fpvA* Δ*fpvB* mutant was 8 μg/mL (Figure 2B), but unexpectedly, DSX also inhibited the growth of the Δ*fpvA* mutant, with the same MIC. This result was surprising, since the Δ*fpvA* mutant has WT susceptibility to TS, suggesting that FpvB is expressed in that background. To confirm that FpvB was expressed in the Δ*fpvA* mutant, we tagged FpvB chromosomally with a C-terminal FLAG tag in both WT and the Δ*fpvA* mutant and blotted for expression. FpvB was detected in the tagged WT and Δ*fpvA* mutant but not in the untagged strains (Figure 2 – figure supplement 1). As a loading control, we monitored expression of PilF, an outer membrane lipoprotein required for multimerization and localization of the *P. aeruginosa* type IV pilus secretin (Koo et al., 2008). Taken together, these results suggested that the role of FpvB in pyoverdine transport needed to be revisited.

**Figure 2.**
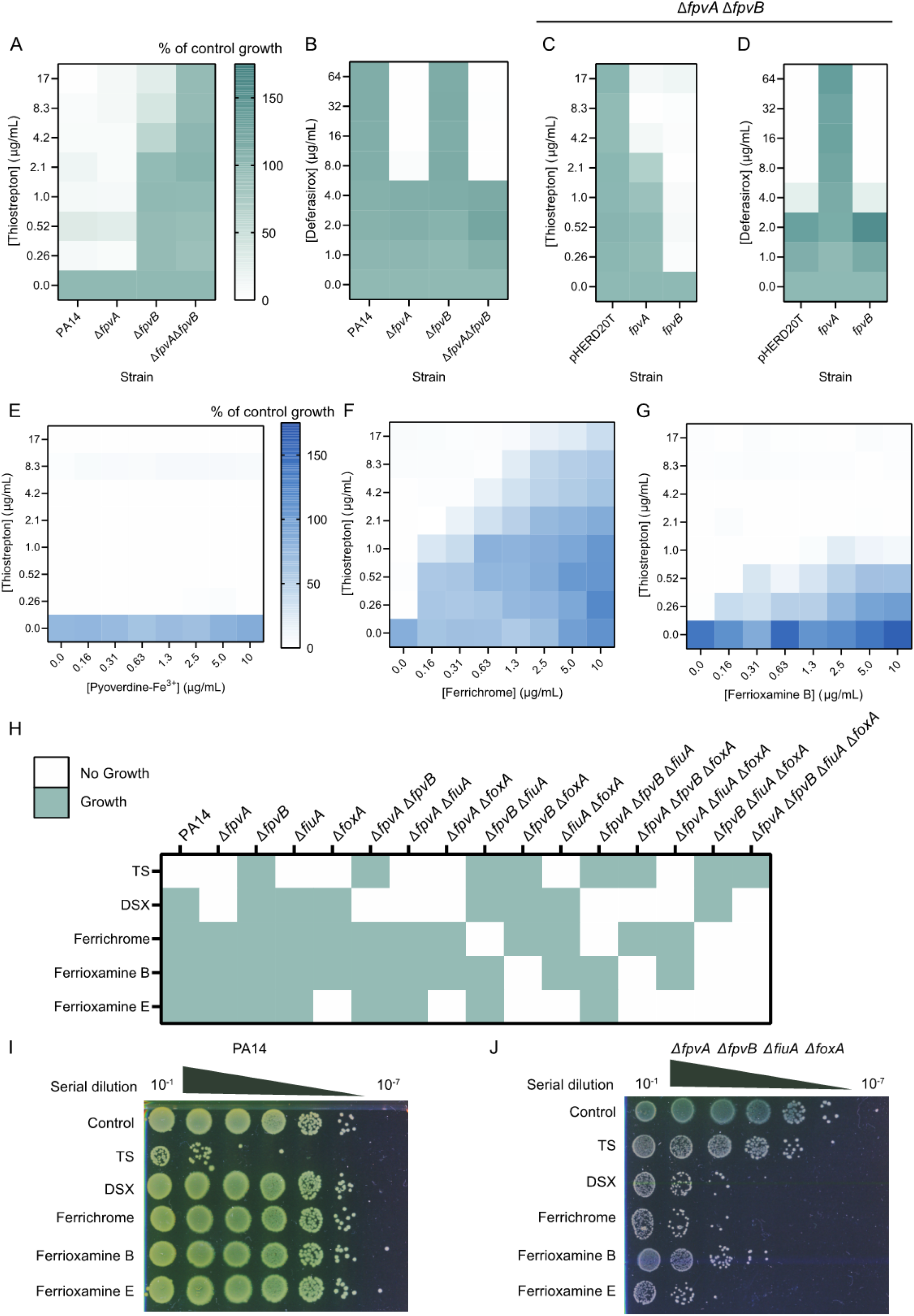
(with 3 supplements). FpvB is required for growth with ferrichrome and ferrioxamine B in the absence of FiuA and FoxA. PA14, Δ*fpvA*, Δ*fpvB*, and Δ*fpvA* Δ*fpvB* treated with (A) TS and (B) DSX in 10:90. Green indicates growth and white indicates lack of growth. Growth is expressed as percent of control. PA14 Δ*fpvA* Δ*fpvB* complemented with pHERD20T, *fpvA*, or *fpvB* treated with (C) TS and (D) DSX in 10:90. Checkerboard assays of PA14 Δ*fpvA* Δ*fpvB* pHERD20T-*fpvB* treated with TS and (E) pyoverdine-Fe^3+^, (F) ferrichrome, and (G) ferrioxamine B. All experiments are averaged from three independent biological replicates. Blue indicates growth and white indicates no growth. (H) Summary of growth phenotypes of WT PA14 and combinations of single, double, triple, and quadruple deletion mutants in *fpvA, fpvB, fiuA*, and *foxA*. Strains were treated with 17 μg/mL TS, 64 μg/mL DSX, 10 μg/mL ferrichrome, 10 μg/mL ferrioxamine B, or 10 μg/mL ferrioxamine E in 10:90. Green indicates growth whereas no growth is indicated in white. Serial 10-fold dilutions of PA14 (I) and the quadruple mutant (J) treated with 17 μg/mL TS, 64 μg/mL DSX, 10 μg/mL ferrichrome, 10 μg/mL ferrioxamine B, or 10 μg/mL ferrioxamine E in 10:90 spotted onto LB 1.5% agar plates and grown overnight at 37°C. Representative plates are shown.

Another explanation for the susceptibility of the Δ*fpvA* mutant to DSX is that the mutant produces less pyoverdine compared to WT cells due to loss of the signalling cascade that controls siderophore production in response to its binding to FpvA (Lamont et al., 2002). To differentiate whether the *fpvA* mutant was susceptible to DSX because of reduced pyoverdine production or because FpvB was a poor pyoverdine transporter, we treated WT, Δ*fpvA*, Δ*fpvB*, and Δ*fpvA* Δ*fpvB* with DSX (64 μg/mL), without and with exogenous pyoverdine (10 μg/mL) for 20 h (Figure 1, Figure 2 – figure supplement 2A). Growth of the Δ*fpvA* mutant should be restored if there are functional pyoverdine transporters. For WT and the Δ*fpvB* mutant, growth was similar between the control and DSX conditions. Pyoverdine or DSX + pyoverdine treatment increased growth. For the Δ*fpvA* mutant, DSX inhibited growth and pyoverdine supplementation delayed growth. DSX + pyoverdine treatment restored growth compared to DSX alone, but also further delayed growth compared to the pyoverdine-alone treatment. The growth of the Δ*fpvA* Δ*fpvB* mutant was inhibited by DSX and pyoverdine, confirming previously published results (Ghysels et al., 2004). Taken together, this suggests that the FpvB is a less efficient pyoverdine transporter than FpvA.

As a control, a Δ*pvdA* Δ*pchA* mutant unable to make pyoverdine and pyochelin was also tested. This mutant was susceptible to DSX. We showed previously that a *pvdA* mutant remains susceptible to TS, indicating that it makes functional pyoverdine transporters even though the signalling cascade controlled by FpvA is disrupted in the absence of ligand production (Ranieri et al., 2019). As expected, pyoverdine restored growth of the mutant in the presence of DSX (Figure 2 – figure supplement 2A). As additional controls, Δ*fpvA*, Δ*fpvA* Δ*fpvB*, and Δ*pvdA* Δ*pchA* were treated with DSX in the presence of the xenosiderophores ferrichrome and ferrioxamine B, which use FiuA and FoxA for uptake (Figure 2 – figure supplement 2B,C) (Llamas et al., 2006). The two xenosiderophores rescued growth of all three mutants in the presence of DSX within a 20 h incubation period, without the delay in growth seen with pyoverdine. Taken together, these data suggest that FpvB is a poor pyoverdine transporter.

The Δ*fpvA* Δ*fpvB* mutant was complemented *in trans* with pHERD20T-*fpvA* or pHERD20T-*fpvB*. pHERD20T is an arabinose-inducible vector with expression driven by the P_BAD_ promoter under control of AraC (Qiu et al., 2008). Complementation with *fpvA* or *fpvB* and induction with arabinose restored TS susceptibility, although susceptibility was greater with *fpvB* (Figure 2C). However, only complementation with *fpvA* restored growth with DSX (Figure 2D). Since *fpvB* could not restore growth of the double mutant in the presence of DSX, we hypothesized that FpvB is a poor transporter of pyoverdine but may transport other siderophores.

### FpvB is a transporter for ferrichrome and ferrioxamine B

We investigated this hypothesis by treating the Δ*fpvA* Δ*fpvB* mutant complemented with *fpvA* or *fpvB* with TS and pyoverdine-Fe^3+^. If pyoverdine competes with TS for the same binding site in the transporter, a reduction in TS susceptibility would be expected as competition would decrease entry of the antibiotic into the cell. TS susceptibility decreased >8-fold with increasing concentrations of pyoverdine-Fe^3+^ when FpvA was expressed (Figure 2 – figure supplement 3A). However, pyoverdine-Fe^3+^ did not impact TS susceptibility when FpvB was expressed under the same conditions (Figure 2E). These results show that pyoverdine is a poor competitor for FpvB binding. As controls, we tested TS with ferrichrome, ferrioxamine B, enterobactin, ferrioxamine E, and arthrobactin, siderophores not expected to use FpvA or FpvB for uptake (Figure 2 – figure supplement 3B-F). None of those siderophores reduced TS susceptibility when FpvA was expressed, showing that only pyoverdine competes with TS for FpvA binding.

Surprisingly, ferrichrome and ferrioxamine B, but not other siderophores, antagonized TS susceptibility when FpvB was expressed (Figure 2F,G, Figure 2 – figure supplement 3G-I). These data suggested that FpvB may transport these two xenosiderophores. We generated 15 single, double, triple, and quadruple knockout mutants lacking different combinations of *fpvA, fpvB, foxA*, and *fiuA* and assessed their growth in the presence of 64 μg/mL DSX, 10 μg/mL ferrichrome, 10 μg/mL ferrioxamine B, and 10 μg/mL ferrioxamine E to determine ligand specificity (Figure 2H). If there were no transporters for a particular siderophore, the mutant would fail to grow due to iron restriction. TS was used as a control at 17 μg/mL since Δ*fpvB* and Δ*fpvA* Δ*fpvB* mutants are resistant to the antibiotic (Figure 2A). Δ*fpvA* mutants were unable to grow in the presence of DSX, consistent with previous results (Figure 1B, Figure 2 – figure supplement 2A). Growth of any mutant combination that included Δ*foxA* was inhibited by ferrioxamine E, confirming previous work showing that ferrioxamine E exclusively uses FoxA as a transporter in *P. aeruginosa* (Normant et al., 2020). Δ*fpvB* Δ*foxA* mutants failed to grow in the presence of ferrioxamine B, while Δ*fpvB* Δ*fiuA* mutants failed to grow with ferrichrome. The quadruple mutant, Δ*fpvA* Δ*fpvB* Δ*fiuA* Δ*foxA*, failed to grow with DSX, ferrioxamine B, ferrioxamine E, or ferrichrome and was resistant to TS. These results suggest that ferrichrome can be taken up via FpvB and FiuA, whereas ferrioxamine B can be taken up via FpvB and FoxA. The siderophores and chelators were bacteriostatic rather than bactericidal, as serial dilution onto non-selective media of PA14 and Δ*fpvA* Δ*fpvB* Δ*fiuA* Δ*foxA* after treatment with the different compounds resulted in regrowth (Figure 2I,J).

### FpvB has higher affinity for ferrichrome and ferrioxamine B than pyoverdine

To determine the affinity of the siderophores for FpvA and FpvB, we adapted the method from Chakravorty et al., to generate a whole-cell sensor where fluorescence quenching serves as an indicator of ligand interaction (Chakravorty et al., 2019). Siderophore binding triggers conformational changes at the extracellular loops of the TBDT (Figure 3A). Cys substitutions were introduced at the loops and labeled with a fluorescent maleimide dye. When a ligand binds the transporter, the loops fold inwards towards the lumen of the barrel, and changes in the chemical environment surrounding the fluorophore lead to quenching (Chakravorty et al., 2019; Kumar et al., 2022). Fluorescence recovery occurs once the siderophore is taken up and the loop returns to its original conformation.

**Figure 3.**
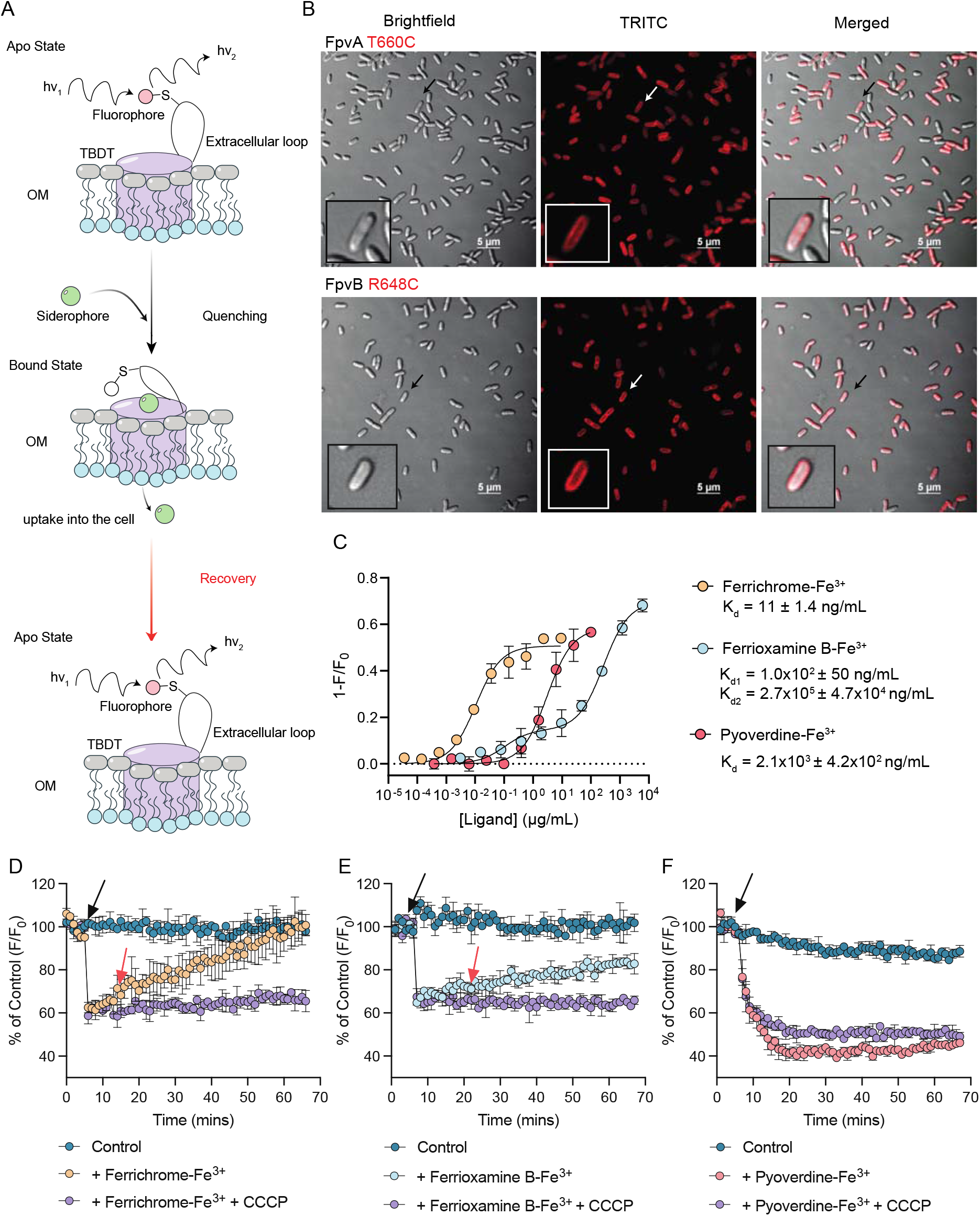
(with 5 supplements). FpvB interacts with ferrichrome, ferrioxamine B, and pyoverdine. (A) Schematic for fluorescence quenching of site-directed labeling of TBDT Cys mutants. A Cys residue is introduced in the extracellular loops of a TBDT of interest and labeled with maleimide dye. A siderophore recognized by the TBDT binds the transporter, inducing conformational changes at the labeled extracellular loop. Changes in the chemical environment surrounding the fluorophore quenches fluorescence. Uptake of the siderophore into the cell restores fluorescence as the loop returns to its original conformation. (B) Fluorescence microscopy images of PA14 Δ*fpvA* Δ*fpvB* pHERD20T-*fpvA* T660C and *fpvB* R684C labeled with AlexaFluor 594. Inset: zoomed in view of a single labeled cell. Scale bar is 5 μm. (C) Fluorescence quenching of labeled FpvB R648C by ferrichrome-Fe^3+^ (blue circles), ferrioxamine B-Fe^3+^ (red circles), or pyoverdine-Fe^3+^ (orange circles). K_d_ values are shown for ferrichrome-Fe^3+^ and pyoverdine-Fe^3+^. Fluorescence recovery of labeled FpvB R648C at ½ K_d_ for (D) ferrichrome-Fe^3+^ (orange circles), (E) ferrioxamine B-Fe^3+^ (blue circles), and (F) pyoverdine-Fe^3+^ (red circles). Teal circles represent vehicle controls and purple circles represent siderophores + 20 μM CCCP. The black arrow indicates when each siderophore complex was added. The red arrow highlights when recovery is observed. No recovery was observed at any pyoverdine-Fe^3+^ concentration. All results are averaged from three independent biological replicates except for the microscopy images where a representative image is shown.

FpvA T660C was used previously to measure pyoverdine-Fe^3+^ binding affinity (Kumar et al., 2022). However, this technique has not been applied to FpvB. A high-confidence structural model of FpvB was generated using AlphaFold2 (Jumper et al., 2021; Mirdita et al., 2022) and aligned with the structure of FpvA (PDB:2O5P) to identify a residue suitable for labeling, with the assumption that FpvA and FpvB undergo similar conformational changes upon ferrisiderophore binding (Figure 3 – figure supplement 1). Based on this analysis, FpvB R648 was mutated to Cys. FpvA T660C and FpvB R648C were each expressed *in trans* in the Δ*fpvA* Δ*fpvB* mutant. FpvB R648C had WT susceptibility to TS and growth in the presence of DSX (Figure 3 – figure supplement 2). Expression of FpvA T660C increased susceptibility to TS by 4-fold compared to WT FpvA and restored growth with DSX. We could also detect expression by fluorescence microscopy, with peripheral labeling consistent with expression in the outer membrane (Figure 3B). Labeling was undetectable in empty vector and WT controls (Figure 3 – figure supplement 3).

The estimated K_d_ of pyoverdine-Fe^3+^ for FpvB was 200-fold higher than FpvA (2.1×10^3^ ± 4.2×10^2^ ng/mL or 1.7×10^3^ ± 3.4×10^2^ nM), confirming that FpvA has higher affinity for pyoverdine (Figure 3C). Ferrichrome-Fe^3+^ and ferrioxamine B-Fe^3+^ also quenched the fluorescence of cells expressing FpvB R648C (Figure 3C). The K_d_ of ferrichrome-Fe^3+^ for FpvB was 11 ± 1.4 ng/mL (15 ± 1.9 nM). Titration of ferrioxamine B-Fe^3+^ yielded a curve with two quenching events, suggesting that it may bind at two sites on FpvB – one of higher affinity than the other. For K_d1_ the binding constant was 1.0×10^2^ ± 50 ng/mL and K_d2_ was 2.7×10^5^ ± 4.7×10^4^ ng/mL. These results may explain the pattern of antagonism between the siderophores and TS. Ferrichrome, which has the highest affinity for FpvB, also had the greatest impact on TS susceptibility. The K_d1_ of ferrioxamine B is approximately 10-fold greater than ferrichrome, suggesting reduced affinity for FpvB, but was sufficient to antagonize TS susceptibility. However, the K_d_ of pyoverdine for FpvB is 200 and 20-fold greater than ferrichrome and ferrioxamine B respectively, and it had the least impact on TS susceptibility.

Siderophore uptake by FpvB was monitored by fluorescence recovery at a concentration of ½ K_d_. In the case of ferrioxamine B, we chose ½ K_d2_ since the quenching signal was greater. Cells were equilibrated for 5 min prior to introduction of the siderophores. Fluorescence was recorded every min for 1 h. Carbonyl cyanide m-chlorophenylhydrazone (CCCP) was used as a control to inhibit fluorescence recovery through dissipation of the proton motive force (PMF) (Klebba et al., 2021; Noinaj et al., 2010), which is required for uptake. Full fluorescence recovery was seen with ferrichrome (Figure 3D). For ferrioxamine B, fluorescence recovered to ~50% of the control; however, no recovery was seen with pyoverdine, suggesting that uptake of this siderophore is slow (Figure 3E,F). These results suggest that of the three siderophores, ferrichrome has the greatest affinity for FpvB and is taken up most rapidly, followed by ferrioxamine B.

### Molecular determinants of TS, ferrichrome, and ferrioxamine B uptake through FpvB

Our data suggested that because ferrichrome, ferrioxamine B, and pyoverdine all quenched fluorescence, they might induce similar conformational changes in FpvB. Ferrichrome-Fe^3+^ and ferrioxamine B-Fe^3+^ were docked into the model of FpvB using AutoDock VINA to identify possible molecular interactions (Jumper et al., 2021; Trott and Olson, 2010). Docking assumed that the siderophore-Fe^3+^ complexes are in similar orientations in their native transporter and in FpvB. Based on the predictions, ferrichrome, and ferrioxamine B bind in a highly aromatic pocket with several Trp and Tyr residues, between the plug and barrel domains (Figure 4A,B).

**Figure 4.**
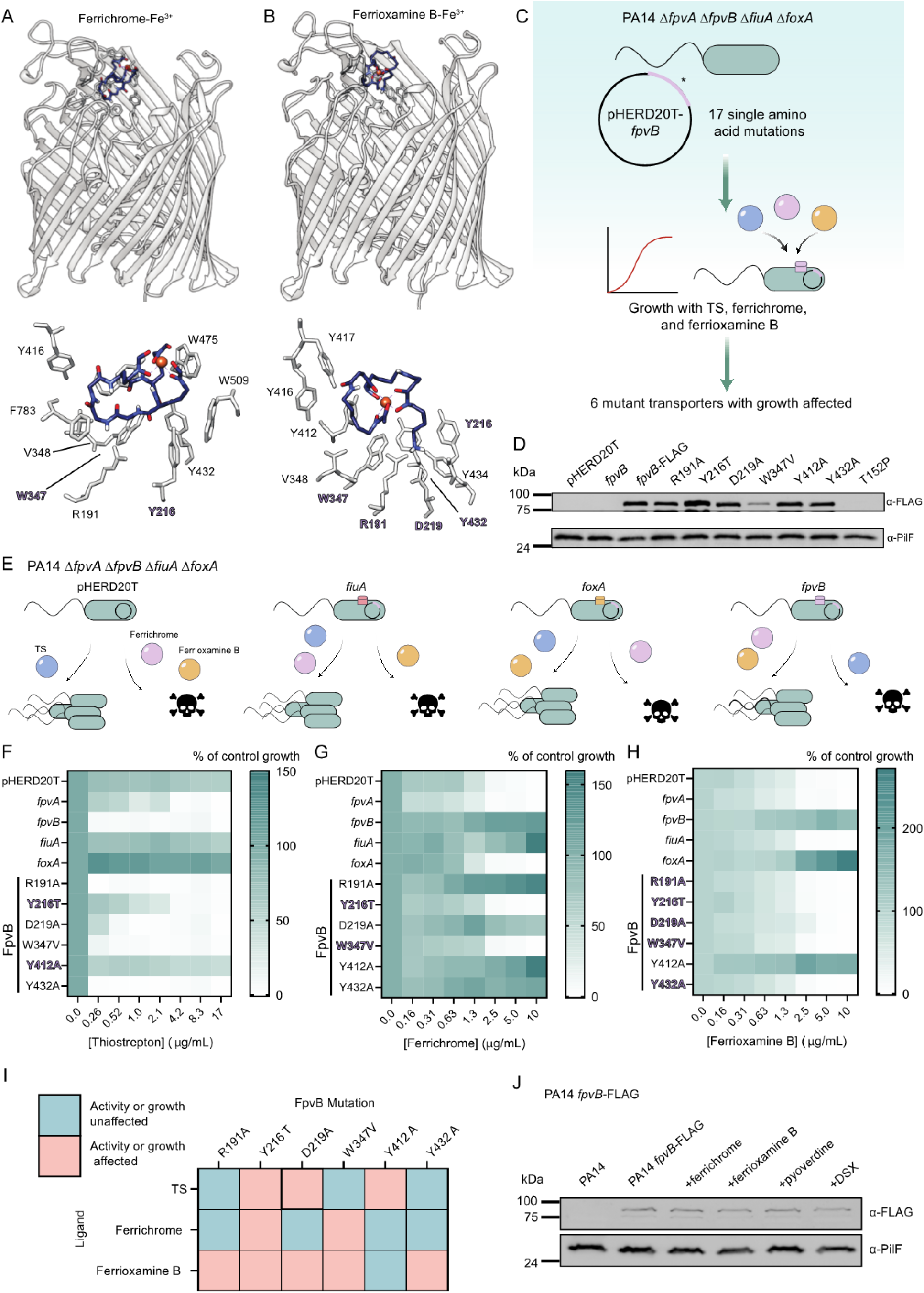
Molecular determinants of TS, ferrichrome, and ferrioxamine B uptake through FpvB. (A) Ferrichrome-Fe^3+^ (blue; PDB: 1BY5) was docked into the AlphaFold2 model of FpvB (grey) using Autodock Vina. Predicted molecular interactions are shown below with residues important for uptake highlighted in purple and bolded. (B) Ferrioxamine B-Fe^3+^ (blue; PDB: 6I96) was docked into the AlphaFold2 predicted model of FpvB (grey). Predicted molecular interactions are shown below with residues important for uptake highlighted in purple and bolded. (C) Schematic for validating the docking predictions. (D) Western blot of FLAG-tagged mutant FpvB transporters, with PilF as a loading control. (E) Expected phenotypes for PA14 Δ*fpvA* Δ*fpvB* Δ*foxA* Δ*fiuA* harbouring pHERD20T, pHERD20T-*fpvB*, pHERD20T-*fiuA*, and pHERD20T-*foxA* treated with TS, ferrichrome, and ferrioxamine B. PA14 Δ*fpvA* Δ*fpvB* Δ*foxA* Δ*fiuA* complemented with empty vector, *fpvA, fpvB, fiuA, foxA* and *fpvB* mutants treated with (F) TS, (G) ferrichrome, and (H) ferrioxamine B. Growth (green) is expressed as percent of control. Results are averaged from three independent biological replicates. (I) Summary of effects of the mutations on TS susceptibility and growth in the presence of ferrichrome and ferrioxamine B. Teal: growth is affected; coral: growth is unaffected. (J) Western blot for WT PA14 with chromosomally integrated C-terminal FLAG tagged FpvB treated with, 10 μg/mL ferrichrome, 10 μg/mL ferrioxamine B, 10 μg/mL pyoverdine, and 64 μg/mL DSX. PilF was used as a loading control for outer membrane proteins.

Site-directed mutagenesis was used to confirm the docking predictions and to define the molecular determinants for ligand uptake through FpvB (Figure 4C). We generated 17 single amino acid substitutions in FpvB and expressed each of them *in trans* in the quadruple Δ*fpvA* Δ*fpvB* Δ*fiuA* Δ*foxA* mutant. Their ability to complement growth of the mutant in the presence of ferrichrome and ferrioxamine B and to restore TS susceptibility was assessed. Of the 17 mutations, six affected TS susceptibility and growth with ferrichrome and ferrioxamine B: R191A (plug domain), Y216T (extracellular loop), D219A (plug domain), W347V (extracellular loop), Y412A (barrel domain), and Y432A (barrel domain). The mutant transporters were tagged with a C-terminal FLAG tag to evaluate expression levels. FLAG-tagged FpvB mutants had WT levels of expression except for Y216T and W347V which were expressed at 140% and 25% of WT respectively. These results suggest that except for W347V, differences in growth are not due to differences in expression (Figure 4D).

The Δ*fpvA* Δ*fpvB* Δ*fiuA* Δ*foxA* mutant was complemented with WT *fpvA*, *fpvB*, *fiuA*, and *foxA in trans* as controls (Figure 4E). The quadruple mutant with the empty vector is resistant to TS and its growth inhibited by ferrichrome and ferrioxamine B. Complementation with *fpvA* was predicted to restore susceptibility to TS but not growth with ferrichrome or ferrioxamine B, while complementation with *fpvB* was expected to restore susceptibility to TS and growth with ferrichrome and ferrioxamine B. Complementation with *fiuA* was expected to restore growth with ferrichrome but not ferrioxamine B, while cells remain resistant to TS. Finally, complementation with *foxA* was expected to restore growth with ferrioxamine B but not ferrichrome, while cells remain resistant to TS. Several mutations within the binding pocket negatively affected TS susceptibility or growth with ferrichrome and ferrioxamine B (Figure 4F-H). Y216T, D219A, and Y412A reduced TS susceptibility. Y216T and W347V prevented growth in the presence of ferrichrome. All mutations except Y412A prevented growth in the presence of ferrioxamine B. The effects of the mutations on TS susceptibility and growth in the presence of ferrichrome and ferrioxamine B are summarized in Figure 4I. Finally, we tested whether ferrichrome, ferrioxamine B, and pyoverdine could stimulate FpvB expression (Figure 4J). DSX was included as a negative control since it does not bind FpvB. None of the compounds stimulated expression, consistent with previous proteomic and RT-PCR assays (Normant et al., 2020).

To determine if these mutations affected binding of ferrichrome and ferrioxamine B to FpvB, we introduced R648C to allow fluorescent labeling of the six point mutants; however, only R191A and W347V could be labeled with AlexaFluor 594 (Figure 5A). Quenching assays with the two xenosiderophores were repeated for the quadruple TBDT mutant expressing WT FpvB, FpvB R191A R648C, or W347V R648C. The R191A mutation increased the K_d_ of ferrichrome-Fe^3+^ by 76-fold to 6.5×10^2^ ± 1.5×10^2^ ng/mL (8.7×10^2^ ± 2.0×10^2^ nM) while the W347V mutation increased the K_d_ >76-fold (Figure 5B). FpvB R191A allowed growth of the quadruple mutant in the presence of ferrichrome whereas W347V did not (Figure 4G,H). These data suggest that certain mutations in FpvB can be tolerated and allow sufficient ferrichrome uptake, potentially because the K_d_ of ferrichrome for WT FpvB is naturally low.

**Figure 5.**
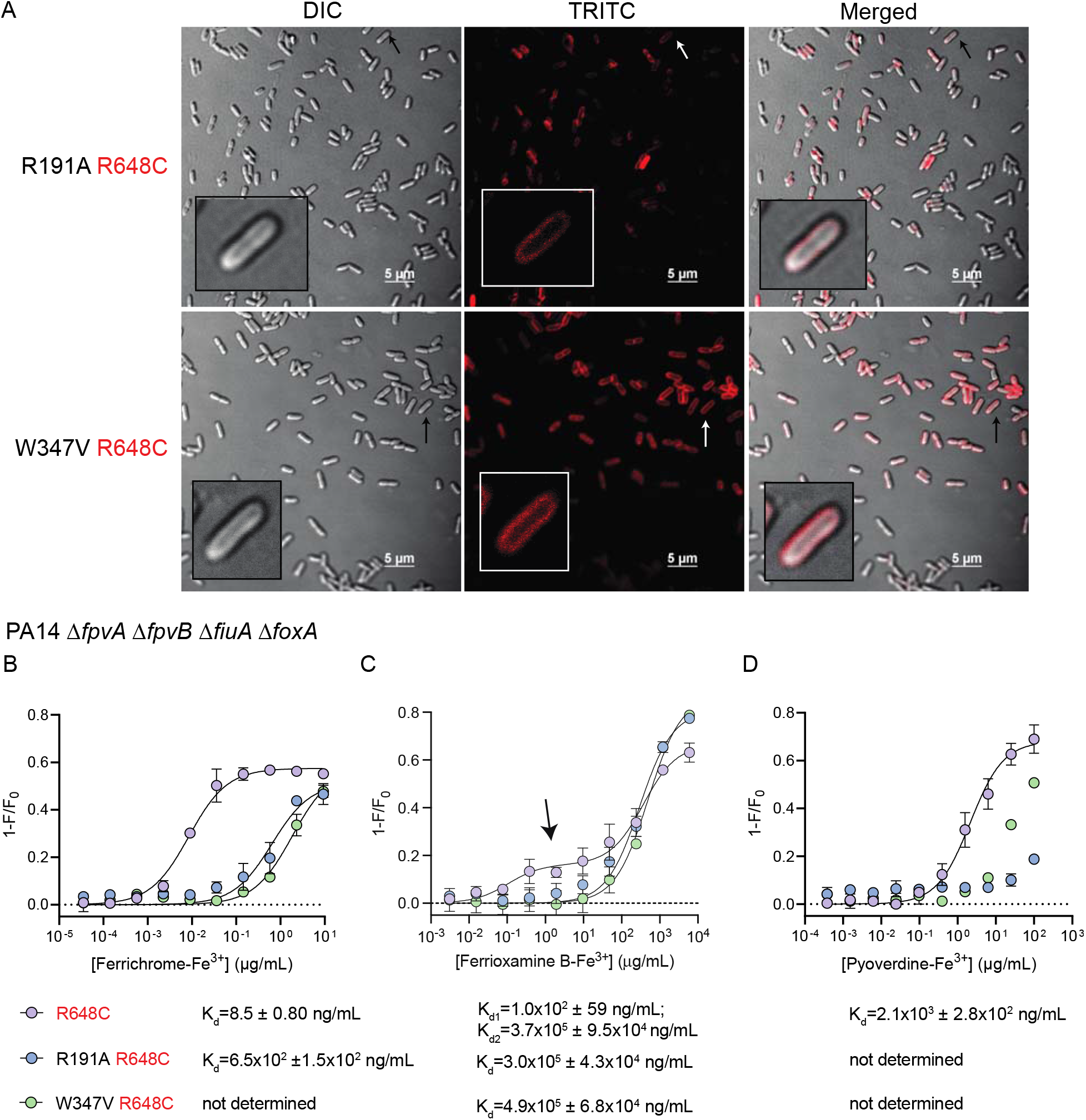
(with 1 supplement). FpvB single residue mutants are stably expressed but have reduced affinity for siderophores. (A) Microscope images of Δ*fpvA ΔfpvB ΔfiuA ΔfoxA* expressing FpvB R191A R648C or W347V R648C labeled with AlexaFluor594. Scale bar: 5 μm. Quenching curves of PA14 Δ*fpvA ΔfpvB ΔfiuA ΔfoxA* expressing fluorescein-5-maleimide-labeled FpvB R648C (purple), R191A R648C (blue), and W347V R648C (green) titrated with (B) ferrichrome-Fe^3+^, (C) ferrioxamine B-Fe^3+^, or (D) pyoverdine-Fe^3+^. The arrow in panel B highlights the first saturation event observed with FpvB R648C titrated with ferrioxamine B-Fe^3+^. Results are averaged from three independent biological replicates.

R191A and W347V also compromised ferrioxamine B uptake, suggesting reduced affinity (Figure 4H,I). This was confirmed with the fluorescence quenching assay using the quadruple transporter mutant expressing FpvB R648C (Figure 5C). The K_d1_ and K_d2_ were 1.0×10^2^ ± 59 ng/mL (1.7×10^2^ ± 96 nM) and 3.7×10^5^ ± 9.5 x10^4^ ng/mL (6.0×10^5^ ± 1.5×10^4^ nM), similar to Δ*fpvA ΔfpvB* (Figure 3C, Figure 5C). The quenching curves for the R191A and W347V mutants fit a typical one-site model rather than the two-site model observed for the WT with K_d_ values of 3.0×10^5^ ± 4.3×10^4^ ng/mL (4.8×10^5^ ± 7.0×10^4^ nM) and 4.9×10^5^ ± 6.8×10^4^ ng/mL (8.0×10^2^ ± 1.1×10^2^ nM) respectively. Neither mutation affected the K_d2_. These results suggest that mutating the predicted binding pocket prevents the initial interaction of ferrioxamine B with FpvB and further supports the hypothesis that ferrioxamine B binds at two distinct sites. Additionally, conformational changes that occur from the first binding event appear to be independent from the second since the quenching is still observed in the mutants.

Pyoverdine-Fe^3+^ binding to FpvB single residue mutants was also assessed using the fluorescence quenching assay. The K_d_s of pyoverdine for FpvB were similar in the Δ*fpvA ΔfpvB* mutant and the quadruple mutant (Figure 3C, Figure 5D). For the R191A and W347V mutants, the K_d_ could not be determined because saturation was not reached even at the highest concentration tested, indicating that pyoverdine has reduced affinity for the two FpvB mutants. Overall, the R191A and W347V mutations reduced the affinity of FpvB for all three ligands.

Since only a subset of FpvB mutant transporters could be fluorescently labeled, we tested competition between TS and ferrichrome via checkerboard assays. The quadruple transporter mutant complemented with WT FpvB or FpvB single-residue mutants was treated with ferrichrome and TS and assessed for antagonism of TS susceptibility (Figure 5 – figure supplement 1). Ferrioxamine B was not tested because the quadruple mutant expressing 5 of 6 FpvB single residue mutants was unable to grow with that siderophore (Figure 4H,I).

Antagonism was observed between ferrichrome and TS when WT FpvB was expressed. Consistent with the data in Figure 4G, the Y216T and W347V mutant FpvB transporters were unable to support growth with ferrichrome. Interestingly, while R191A and D219A supported growth of the quadruple mutant in the presence of ferrichrome, no competition with TS was observed, suggesting that these mutations specifically reduce the affinity of FpvB for ferrichrome. When the Y432A mutant transporter was expressed, ferrichrome antagonised TS activity, although to a lesser extent than observed with the WT. Overall, the mutations diminished the ability of ferrichrome to compete with TS.

To identify any other proteins important for susceptibility to TS, we raised mutants resistant to TS + DSX. PA14 was grown in liquid cultures with 17 μg/mL of TS + 64 μg/mL DSX and passaged over a period of three weeks. Single colonies were isolated on agar containing 17 μg/mL of TS + 64 μg/mL DSX and 6 were selected for sequencing. Two mutations that conferred resistance were identified. All resistant mutants had a premature stop codon in *tonB1* after residue A36, suggesting that functional TonB1 is not made. We showed previously that *tonB1* mutants are resistant to thiopeptides (Chan and Burrows, 2021). A mutation in *fpvB* (T152P) was also identified in two colonies. FpvB T152P was tagged with a C-terminal FLAG tag and expressed *in trans* to evaluate expression but no detectable bands were present from outer membrane preparations (Figure 4D). Therefore, this mutation likely abolishes expression of functional FpvB. No mutations in FpvA were identified, although this may be due to selective pressure from DSX since we showed that FpvA is important for growth with the chelator (Figure 2B,D).

## DISCUSSION

This work expands on the discovery of FpvB as an alternative transporter for pyoverdine (Ghysels et al., 2004). Ghysels et al. made a *P. aeruginosa* PAO1 mutant unable to produce FpvA, pyoverdine, or pyochelin, which grew in casamino acids (CAA) medium. Supplementing the media with ethylene diamine di(o-hydroxyphenyl)acetic acid (EDDHA), an iron chelator unable to enter cells, inhibited growth, similar to our results with the PA14 Δ*fpvA* mutant and DSX (Figure 2 – Figure Supplement 1). Supplementing the media with pyoverdine restored growth of the mutant after 24 h but not 12 h, suggesting that there is a TBDT with limited affinity for pyoverdine. Deleting *fpvB* inhibited growth recovery, suggesting that FpvB transports pyoverdine. Another study showed that a PAO1 Δ*fpvA* mutant was unable to grow in media supplemented with EDDHA even if pyoverdine was added, although growth was only monitored for 10 h rather than 24 h (Shen et al., 2002). Similarly, a PAO1 FpvA-deficient strain was unable to take up iron from pyoverdine-^59^Fe, regardless of whether pyoverdine from PAO1 or other Pseudomonads was provided, although uptake was measured only for 15 min (Meyer et al., 1999). Together, these results suggest that FpvB is a poor transporter for pyoverdine, which informed our hypothesis that FpvB may transport other siderophores.

In this work, we uncovered additional uptake pathways for the fungal siderophore, ferrichrome, and the bacterial siderophore, ferrioxamine B, in *P. aeruginosa*. Canonically, ferrichrome is taken up via FiuA while ferrioxamine B uses FoxA. Both siderophores can be recognized by FpvB (Figure 6), and ferrichrome binds with greater affinity than ferrioxamine B or pyoverdine, based on fluorescence quenching data (Figure 3C). Unexpectedly, the quenching curve of ferrioxamine B-Fe^3+^ supported a two-site binding model. This result suggests that the binding mode of ferrioxamine B for FpvB is different than those of ferrichrome and pyoverdine, and shows that different ligands can interact in distinct ways with the same TBDT. A two-site model has been proposed for binding of enterobactin to its transporter PfeA, but ferrioxamine B and E were reported to bind their primary transporter FoxA only at a single site (Josts et al., 2019; Moynié et al., 2019; Normant et al., 2020). We also used fluorescence recovery as an indicator of ligand uptake. Fluorescence recovery was not observed for pyoverdine even after 1 h (Figure 3F), suggesting that pyoverdine uptake through FpvB is slow. A combination of a high K_d_ and slow uptake may explain why FpvB is a poor transporter for pyoverdine.

**Figure 6.**
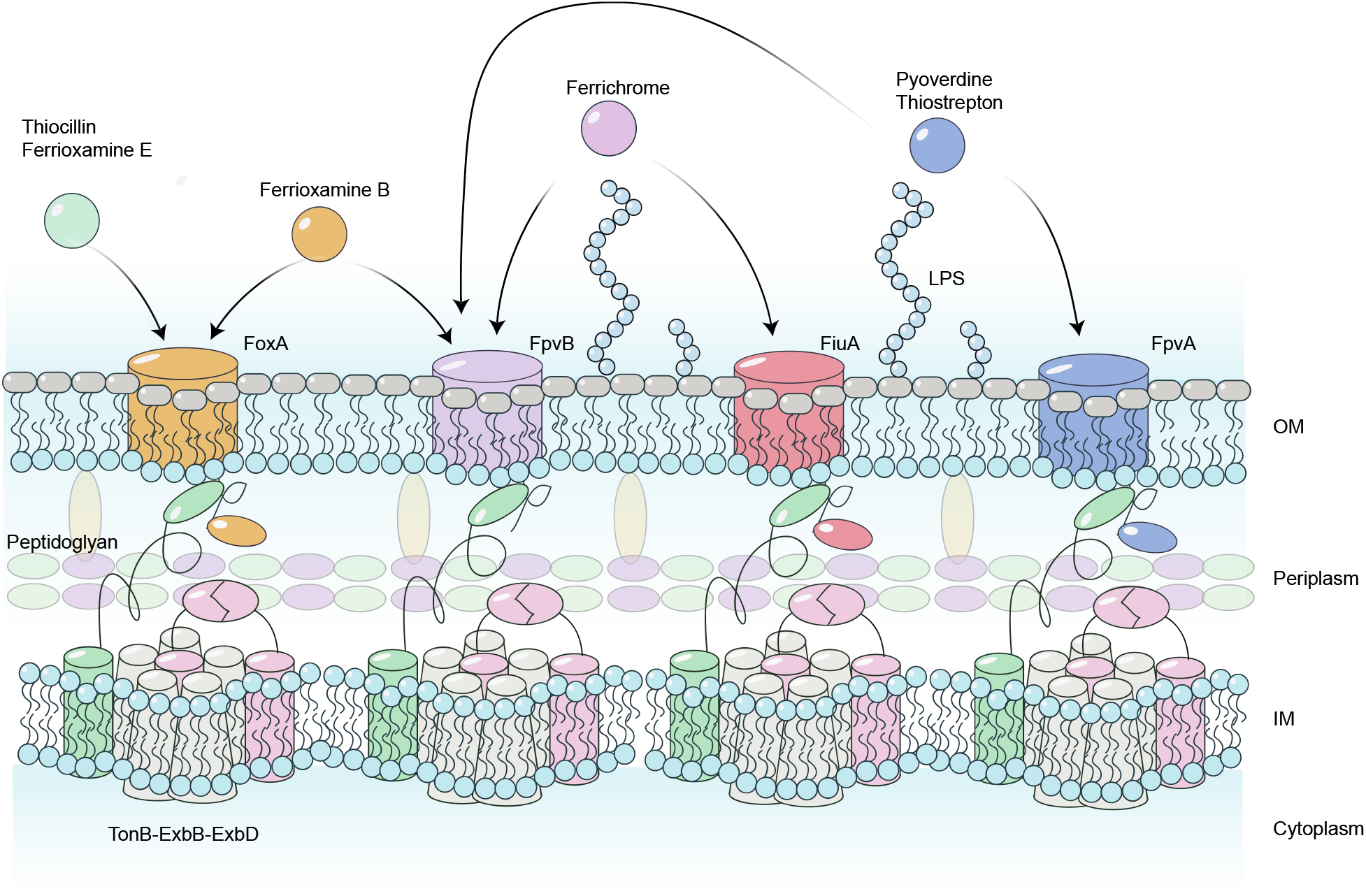
Model for thiopeptide and siderophore uptake through the TBDTs FpvA, FpvB, FoxA, and FiuA in *P. aeruginosa*. OM – outer membrane; IM – inner membrane.

The site-directed mutagenesis data suggest that all three ligands bind a similar hydrophobic pocket in FpvB, but the molecular determinants for uptake depend on the ligand. For example, W347 is important for both ferrichrome and ferrioxamine B but not TS uptake. Y412 is important for TS uptake but not for the two xenosiderophores, and Y216 is essential for uptake of all three ligands. Co-structures of FpvB with each of ligands will be informative.

Combining the site-directed mutagenesis data with fluorescence quenching and checkerboard competition assays, we showed that modifying the hydrophobic pocket of FpvB reduced its affinity for the ligands. With ferrioxamine B, the R191A and W347V mutations abolished the initial quenching event observed in WT FpvB (Figure 5C). However, the second quenching event was unaffected, suggesting that the two binding events occur independently, and that conformational changes caused by the first event are not required for the second. Future work will focus on locating the second site, to understand how ferrioxamine B interacts with FpvB. Additionally, this work supports the idea that the binding mechanism of different siderophores can vary even for the same transporter.

Determining the ligand specificity of TBDTs can be challenging. Traditionally, sequence alignments of transporters with known ligands and transporters with unknown ligands were used to make inferences about the function of the new transporter. This method works well for those that share high sequence similarity. For example, the aerobactin transporter in *P. aeruginosa* was discovered by comparing the sequences of *Escherichia coli* IutA and *P. aeruginosa* ChtA which share 46% identity (63% similarity) (Cuív et al., 2006). FpvB was initially discovered as a secondary ferripyoverdine transporter using this method, as it shares 54% amino acid similarity with FpvA (Ghysels et al., 2004). However, alignments do not provide a comprehensive picture of the range of ligands that can be taken up. FpvB shares only 30-40% similarity with FoxA and FiuA, even though our data suggest that it has higher affinity for ferrichrome than pyoverdine.

Proteomic and RT-PCR approaches have also been used to identify the transporters for ferrioxamine E, ferrioxamine B, and ferrichrome in *P. aeruginosa* (Normant et al., 2020). FoxA expression was upregulated in the presence of ferrioxamine E and ferrioxamine B, while FiuA expression was upregulated by ferrichrome. However, only a subset of TBDTs have N-terminal signalling domains that respond to the presence of ferrisiderophores in a feed-forward regulation loop to increase their expression. Further, many TBDTs have redundant functions and knocking out single transporters may be insufficient to abolish uptake, making it difficult to understand the complete uptake pathway for a ligand. As an alternative, competition between an antimicrobial and siderophore has been used to show that an antimicrobial exploits a particular TBDT for uptake (Ferguson et al., 2001; Southwell et al., 2021). Here, we used competition between TS and various siderophores to show that FpvB recognizes ferrichrome and ferrioxamine B, antagonizing TS susceptibility. This method may be applicable to study the range of ligands that can be recognized by TBDTs in bacteria besides *P. aeruginosa*. The disadvantage of this method is that a antimicrobial known to use the TBDT of interest for uptake is required.

Previous studies established that 93% of *P. aeruginosa* clinical and environmental isolates have *fpvB* (Bodilis et al., 2009; Ghysels et al., 2004; Ranieri et al., 2019). It may be advantageous to produce a single transporter that recognizes at least three different siderophores; therefore, the observation that 7% of isolates lack *fpvB* seems paradoxical. Loss of *fpvB* may improve the fitness of *P. aeruginosa* in the lungs of cystic fibrosis patients through genome reduction (Dingemans et al., 2014). Supporting this finding, we previously identified a TS-resistant but thiocillin-sensitive clinical isolate, C0379, missing ~800 bp from the 5’ region of *fpvB*, suggesting that it may have once produced functional FpvB (Ranieri et al., 2019). Δ*fpvB* mutants are also fitter than WT cells when treated with the antibiotic gentamicin (González et al., 2021). Its absence may reduce the metabolic burden on cells living in stressful environments whether due to nutrient limitation or antibiotic stress. However, the proportion of isolates from environmental or clinical sources lacking *fpvB* is similar, suggesting that there are multiple factors involved (Bodilis et al., 2009). For example, while we identified TS + DSX-resistant mutants with a single point mutation in *fpvB* (T152P) that abolished its expression, we also identified resistant *P. aeruginosa* unable to make TonB1.

This work has implications for our understanding of how siderophores are taken up by *P. aeruginosa* compared to other bacteria. In *E. coli*, ferrichrome is recognized by FhuA whereas ferrioxamine B weakly binds FhuE, which does not take up ferrioxamine E or ferrichrome (Grinter and Lithgow, 2019; Locher et al., 1998). In *P. aeruginosa*, ferrichrome is recognized by FiuA and FpvB, ferrioxamine B is recognized by FoxA and FpvB, and ferrioxamine E is exclusively recognized by FoxA. FpvB is unusual in that TBDTs typically take up siderophore-iron complexes that are structurally related to their native siderophores, which suggests that FpvB may have a high degree of promiscuity compared to other TBDTs. These differences in uptake are important considerations in the design of broad-spectrum siderophore-antibiotic conjugates for gram negative pathogens.

There is growing interest in the use of antimicrobials that exploit TBDTs for uptake. For example, conjugating antibiotics to ferrichrome or ferrioxamine B may permit a compound to be taken up by both FpvB and FiuA or FoxA. The benefits of an antibiotic-siderophore conjugate that can use multiple receptors include a reduced chance of developing resistance, as cells would have to lose multiple TBDTs. In this context, it would be of interest to see how modifications to the siderophore structure affects binding and uptake. For example, one natural variation that prevents ferrioxamine E from using FpvB is that it is cyclic (Figure 1) whereas ferrioxamine B is linear in its apo-form; however, they both adopt similar cyclic conformations when bound to Fe^3+^. Further, ferrioxamine B has an amine tail predicted to interact with R191 that does not participate in iron chelation; this extension is absent in ferrioxamine E. Structure-activity relationship studies of ferrichrome and ferrioxamine B may further reveal ligand-transporter interactions to allow for the design of better siderophore-drug conjugates.

## METHODS AND MATERIALS

### Strains and primers

All strains and primers used in this study are listed in the supplemental materials (**Supplementary Table S1**).

### Compounds and media

Ferrichrome and fluorescein-5-maleimide were purchased from Cayman Chemicals. Ferrioxamine B was purchased from Calbiochem. Pyoverdine and pyoverdine-Fe^3+^ was purchased from Sigma. Arthrobactin was purchased from MolPort. AlexaFluor594 C5 maleimide was purchased from Fisher Scientific. Carbenicillin was purchased from AK Scientific. L-arabinose was purchased from Bioshop. Glucose was purchased from Fisher Scientific. Stock solutions and powders were stored at −20°C. Lysogeny broth (LB) was purchased as a premixed powder from Bioshop. 10:90 medium was prepared as previously described (Chan and Burrows, 2021; Ranieri et al., 2019). L-arabinose was prepared as a 20% (wt/v) stock solution in 10:90 and filter sterilized (0.2μm pore size - Fisherbrand).

### Molecular biology

Chromosomal mutants were generated by allelic exchange using pEX18Gm (Hoang et al., 1998). Primers flanking the upstream and downstream regions of each gene of interest were amplified from PA14 genomic DNA (Promega Wizard Genomic DNA Purification Kit) and extracted with GeneJet Gel Extraction Kit (ThermoFisher). The upstream and downstream regions were joined by overlap extension PCR or ligation, digested with the indicated enzymes (FastDigest ThermoFisher), and ligated into pEX18Gm to make each deletion construct (T4 DNA ligase, ThermoFisher) (Hoang et al., 1998). The ligation mixtures were transformed into competent *E. coli* DH5α by heat shock with a recovery period of 2-3 h in LB. Cells were plated on LB 1.5% agar containing 15 μg/mL gentamicin supplemented with 5-bromo-4-chloro-3-indolyl-β-D-galactopyranoside (X-gal) for blue-white screening. The plates were incubated at 37°C overnight. Colony PCR was performed on white colonies to check for the correct inserts and those with the insert were grown in LB + 15 μg/mL gentamicin overnight at 37°C with shaking (200 rpm). Plasmids were isolated using GeneJet Plasmid Miniprep Kit (ThermoFisher).

Plasmids with correct inserts were transformed into competent *E. coli* SM10 by heat shock with a recovery period of 2-3 h in LB. The cells were plated on LB 1.5% agar containing 10 μg/mL gentamicin and grown overnight at 37°C. One colony was picked and inoculated in LB + 10 μg/mL gentamicin. The *P. aeruginosa* mutant of interest was also inoculated from a single colony in LB. Both cultures were grown overnight at 37°C with shaking (200 rpm). SM10 with the desired deletion construct was mated with PA14 by mixing equal volumes of each overnight culture in a 1.5mL centrifuge tube. Cells were spun down and the supernatant was removed. Cells were resuspended in 50 μL fresh LB, spotted on LB 1.5% agar, and incubated overnight at 37°C. Cells from the mating spot were streaked onto *Pseudomonas* Isolation Agar (PIA, Difco) supplemented with 100 μg/mL gentamicin and incubated overnight at 37°C. Single colonies were streaked onto LB (no salt) + 15% sucrose (BioShop) and incubated overnight at 37°C. To check for colonies with the correct deletion, 16 colonies were patched onto LB + 15% sucrose and LB + 30 μg/mL gentamicin and incubated overnight at 37°C. Patches that grew on the sucrose plates but not gentamicin plates were checked by colony PCR with primers flanking the deleted gene and internal primers and compared to WT controls. Patches with the desired deletions were streaked onto LB + 15% sucrose to isolate single colonies, incubated overnight at 37°C and checked again by colony PCR. A single colony was inoculated into LB broth and the process was repeated to generate double, triple, and quadruple mutants.

Complemented strains were made using the plasmid pHERD20T, an arabinose-inducible expression vector with the P_BAD_ promoter under control of AraC (Qiu et al., 2008). Primers flanking the gene of interest including the native ribosome binding site were amplified from *P. aeruginosa* PA14 genomic DNA, digested with the desired restriction enzymes, and ligated into pHERD20T digested with the same enzymes. Ligation mixtures were added to chemically competent DH5α, and DNA introduced by heat shock with a recovery period of 1-2 h in LB at 37°C. All the cells were plated on LB 1.5% agar supplemented 100 μg/mL ampicillin, X-gal, and 0.1% arabinose and incubated at 37°C overnight. White colonies were analyzed by colony PCR and colonies with plasmids containing the desired insert size were cultured in LB broth supplemented with 100 μg/mL ampicillin. Plasmids were isolated, inserts validated by restriction digest, and electroporated into the desired *P. aeruginosa* strain or mutant with a recovery of 1-2 h in LB at 37°C. All the cells were plated on LB 1.5% agar supplemented with 200 μg/mL carbenicillin (AK Scientific). A single colony was picked and grown in LB supplemented with 200 μg/mL carbenicillin overnight at 37°C and used to make glycerol stocks and for subsequent assays. Correct inserts were verified by Sanger sequencing by the McMaster Genomics Facility Mobix Lab.

### MIC assays

MIC assays were conducted as previously described (Chan and Burrows, 2021; Ranieri et al., 2019). Overnight cultures were grown in LB from a glycerol stock at 37°C with shaking (200 rpm). Subcultures in 10:90 (1:100 dilution) were cultured for 4 h. Cells were adjusted to an OD_600_ of 0.1/500 in 10:90. All compounds were serially diluted 2-fold in DMSO or H_2_O at 75x the final concentration. Plates were sealed to prevent evaporation and incubated at 37°C overnight in a shaking incubator (200 rpm). The next day, the OD_600_ was determined with a plate reader (Thermo Scientific) and normalized to percent of growth of the vehicle control after subtracting the OD_600_ from blank media.

### Checkerboard assays

Checkerboards (8 rows x 8 columns) were conducted as previously described in a 96-well plate (Nunc) (Chan and Burrows, 2021; Ranieri et al., 2019). TS at 75x the final concentration dissolved in DMSO was added in columns from bottom to top in increasing concentrations. Siderophores at 75x the final concentration dissolved in DSMO were added from in rows from left to right in increasing concentrations. Four columns were used for vehicle controls (DMSO) and sterile controls. Media with bacteria as described in the MIC assays were added to obtain a final volume of 150 μL. Plates were sealed to prevent evaporation and incubated at 37°C overnight in a shaking incubator (200 rpm). The next day, the OD_600_ was determined with a plate reader (Thermo Scientific) and normalized to percent of growth of the vehicle control (DMSO) after subtracting the OD_600_ from blank media.

### Microscopy, fluorescence quenching, and recovery assay

Cells were cultured overnight in LB with carbenicillin (200 μg/mL) at 37°C with shaking at 200 rpm. Cells were subcultured (1:200 dilution) in 10:90 + 2% arabinose for four hours without carbenicillin at 37°C with shaking at 200 rpm. Cells were harvested by centrifugation for 5 min and 6000 G and resuspended in sterile 1X phosphate buffered saline (PBS; pH 7.4; 1X PBS was made from a 10X stock (80 g NaCl, 2 g KCl, 26.8 g Na_2_ HPO_4_-7H_2_O, 2.4 g KH_2_PO_4_ in 1 L of deionized H_2_O) with 10 μM AlexaFluor594 C_5_-maleimide for microscopy or fluorescein-5-maleimide for fluorescence quenching and recovery assays and incubated in the dark at room temperature for 30 min on a shaking incubator at 37°C (200 rpm). Excess dye was quenched with 1 mM dithiothreitol (DTT) (Sigma) to stop the reaction. Cells were washed 3X with PBS. For fluorescence recovery assays, cells were incubated with 1X PBS + 0.4% glucose at 37°C with shaking at 200 rpm. Glucose was prepared as a 20% stock (wt/v) in 1X PBS and filter sterilized. Cells were spun down and washed 3X with 1X PBS and resuspended in 1X PBS for quenching assays and 1X PBS + 0.4% glucose for fluorescence recovery assays.

For microscopy, AlexaFluor594 C_5_-maleimide-labeled cells were spotted onto a 1% agarose pad on a microscope slide. The agarose pad was mounted with a glass coverslip directly prior to imaging. Cells were imaged using brightfield and fluorescence microscopy on a Nikon A1 confocal microscope through a Plan Apo 60X (NA=1.40) oil objective. Image acquisition was done using Nikon NIS Elements Advanced Research (Version 5.11.01 64-bit) software.

For fluorescence quenching and recovery assays, cells were diluted to an OD_600_ of 0.1 for both types of assays and 148 μL was added into wells of a 96-well black plate (Corning). Fluorescence was recorded for 5 mins at 1 min intervals at 37°C (BioTek Neo; Ex. 494nm; Em. 520nm). After 5 min, 2 μL of each serial dilution for ferrioxamine-Fe^3+^, ferrichrome-Fe^3+^, and pyoverdine-Fe^3+^ or DMSO for vehicle controls was added to each well. Fluorescence was recorded immediately for quenching assays and for 1 h at 1 min interval for fluorescence recovery assays. Background fluorescence was subtracted from all fluorescence readings and 1-F/F_0_ was used to calculate the degree of quenching where F0 is the initial fluorescence. K_d_ was calculated using GraphPad Prism using a one-site specific binding model or a two-site model. For fluorescence recovery assays, background was subtracted from all fluorescence readings and F/F_0_ expressed as a percent of control was plotted versus time.

### Outer-membrane preparations, SDS PAGE, and Western Blots

The quadruple mutant expressing mutant FpvB transporters was grown overnight in LB with 200μg/mL carbenicillin at 37°C with shaking (200rpm). Cells were subcultured 1:100 into 10:90 for four h then diluted to OD_600_ 0.1/500 in 50mL of fresh 10:90 + 2% arabinose. Cultures were grown overnight at 37°C with shaking (200 rpm). Cells were harvested by centrifugation (5 min, 6000 G) and resuspended in 10 mM Tris pH 8.0.

Cells were lysed by sonication (Misonix Sonicator 3000) on ice (30s pulse, power level 5.0). Cell debris was removed by centrifugation (6000 G, 5 min, 4°C). Proteins were harvested at 21000 G for 30 min at 4°C. The pellet was resuspended in 100 μL de-ionized H_2_O and combined with 900 μL 11.1 mM Tris, 1% sarkosyl (Fisher Scientific), pH 7.6 and incubated at room temperature for 30 min with shaking (200 rpm). Outer membrane pellets were collected by centrifugation at 21000 G for 30 min at 4°C.

Outer membrane preparations were resuspended in 20 μL 1X loading buffer for SDS-PAGE analysis. SDS-PAGE buffer was made from a 10X tris-glycine buffer stock (30.3 g tris, 144 g glycine, and 20 mL 10% SDS). Each lane was loaded with 10 μL of outer membrane prep and proteins separated at 80V for 10 min and 120V for 1.5 h. Proteins were transferred to nitrocellulose membranes (225mA, 1 h, in transfer buffer (20% methanol, 100 mL of a 10X trisglycine buffer stock without SDS) and blocked with 5% skim milk in PBS overnight at 4°C. Primary antibodies (mouse α-DYKDDDDK (Invitrogen MA1-91878), and rabbit #3198 α-PilF)) were used at 1:1000 dilutions in PBS and incubated with the blot at room temperature for 1 h. Blots were washed 3×10 min with PBS and incubated with rabbit α-mouse-alkaline phosphatase for 2 h in PBS at 1:2000 dilutions in PBS. Blots were washed with PBS 3x (10 mins per wash) before incubation with alkaline phosphatase buffer (1 mM Tris, 100 mM NaCl, 5 mM MgCl_2_ pH 9.5) + 5-bromo-4-chloro-3-indoyl phosphate (BCIP) + nitro-blue tetrazolium (NBT) for 15-30 min. Blots were developed in the dark and imaged on an Azure C400 Imaging System. Band densities were quantified using ImageJ (Schneider et al., 2012).

### Mutants resistant to TS

PA14 was cultured in 10:90 supplemented with 17 μg/mL TS + 64 μg/mL DSX at 37°C with shaking at 200 rpm (Ranieri et al., 2019). Cells were passaged when turbidity was evident (1:500 dilution) into fresh media with the same concentrations of TS + DSX. This procedure was repeated for 3 weeks until cells grew overnight with TS + DSX. The cells were streaked on to LB deferrated with FEC-1, to remove iron, overnight at 37°C and supplemented with TS + DSX. FEC-1 was supplied by Chelation Partners Inc. (Fe Pharmaceuticals) (Parquet et al., 2018). Isolates were picked, cultured in LB, and chromosomal DNA was isolated and whole genome sequencing performed by the McMaster Genomics Facility Mobix Lab. Breseq was used to identify mutations associated with TS+DSX resistance (Deatherage and Barrick, 2014).

### Structural Comparisons and Docking

Structural models of FpvB were generated using AlphaFold2 (ColabFold) and compared to the crystal structure of FpvA (PDB: 2O5P) (Jumper et al., 2021; Mirdita et al., 2022). Structural comparisons of FpvB and FpvA were visualized using Chimera (Pettersen et al., 2004). Ferrichrome (PDB: 1BY5) and ferrioxamine B (PDB:6I96) were docking into the AlphaFold2 model of FpvB using AutoDock VINA (Trott and Olson, 2010) with a box size of center_x = - 7.729, center_y = 34.958, center_z = 113.797, size_x = 114, size_y = 116, size_z = 118. The top pose generated was used for further studies.

## Acknowledgements

We thank Dr. David Sychantha for his valuable comments on the quenching curve of ferrioxamine B. This work was supported by a Natural Sciences and Engineering Research Council Discovery Grant RGPIN-2021-04237 to LLB. LLB holds a Tier 1 Canada Research Chair in Microbe-Surface Interactions. DCKC holds a Canadian Institute of Health Research (CIHR) Canada Graduate Scholarship – Doctoral program (CGS-D).

---

**Figure 2 – figure supplement 1.**
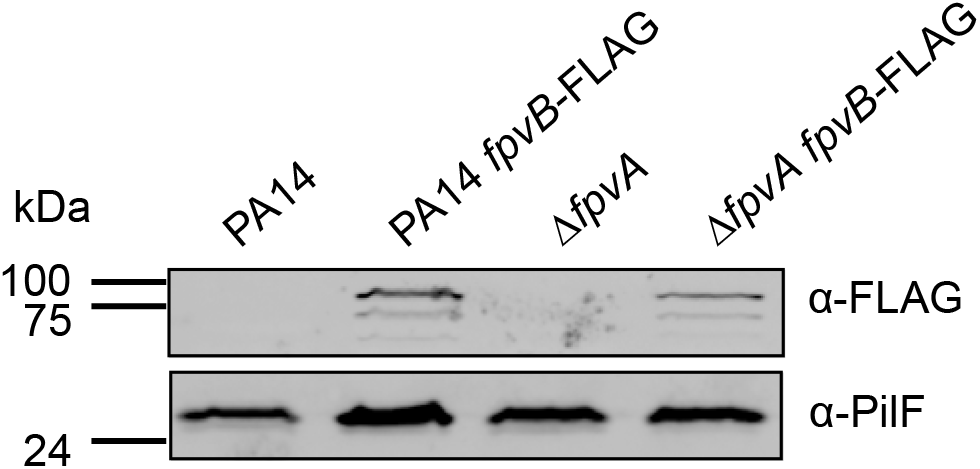
FpvB is expressed in WT PA14 and the Δ*fpvA* mutant. PilF was used as a loading control for outer membrane proteins.

**Figure 2 – figure supplement 2.**
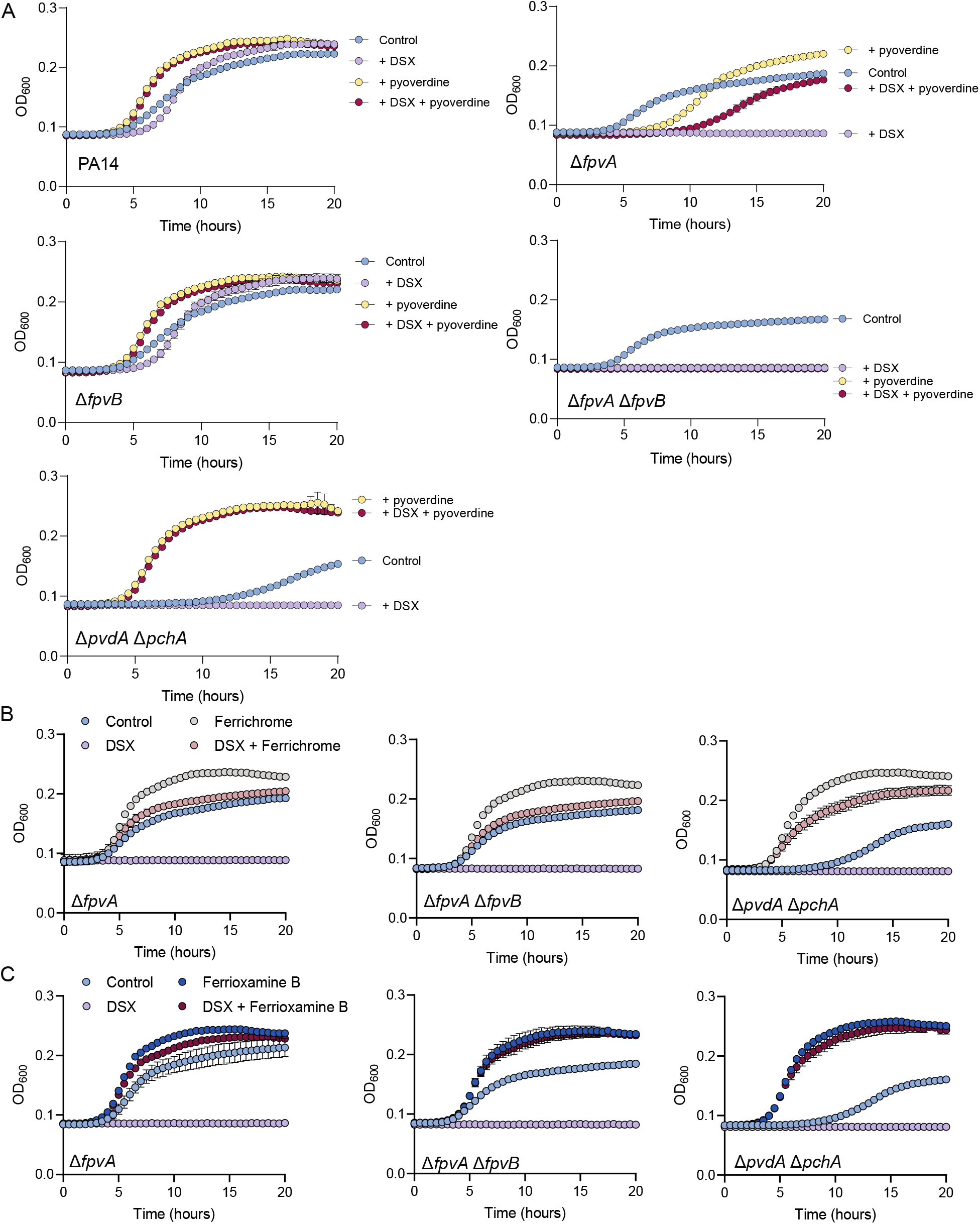
Growth curves of PA14 and mutants with DSX and different siderophores in 10:90. (A) PA14, Δ*fpvA*, Δ*fpvB*, Δ*fpvA* Δ*fpvB*, and Δ*pvdA* Δ*pchA* growth. Blue indicates control, purple indicates treatment with 64 μg/mL DSX, yellow indicates treatment with 10 μg/mL pyoverdine (PA14) and red indicates treatment with 64 μg/mL DSX and 10 μg/mL pyoverdine (PA14). (B) Δ*fpvA*, Δ*fpvA* Δ*fpvB*, and Δ*pvdA* Δ*pchA* growth. Blue indicates control, purple indicates treatment with 64 μg/mL DSX, grey indicates treatment with 10 μg/mL ferrichrome, and red indicates treatment with 64 μg/mL DSX and 10 μg/mL ferrichrome. (C) Δ*fpvA*, Δ*fpvA* Δ*fpvB*, and Δ*pvdA* Δ*pchA* growth. Blue indicates control, purple indicates treatment with 64 μg/mL DSX, grey indicates treatment with 10 μg/mL ferrioxamine B, and red indicates treatment with 64 μg/mL DSX and 10 μg/mL ferrioxamine B. Results are averaged from three independent biological replicates.

**Figure 2 – figure supplement 3.**
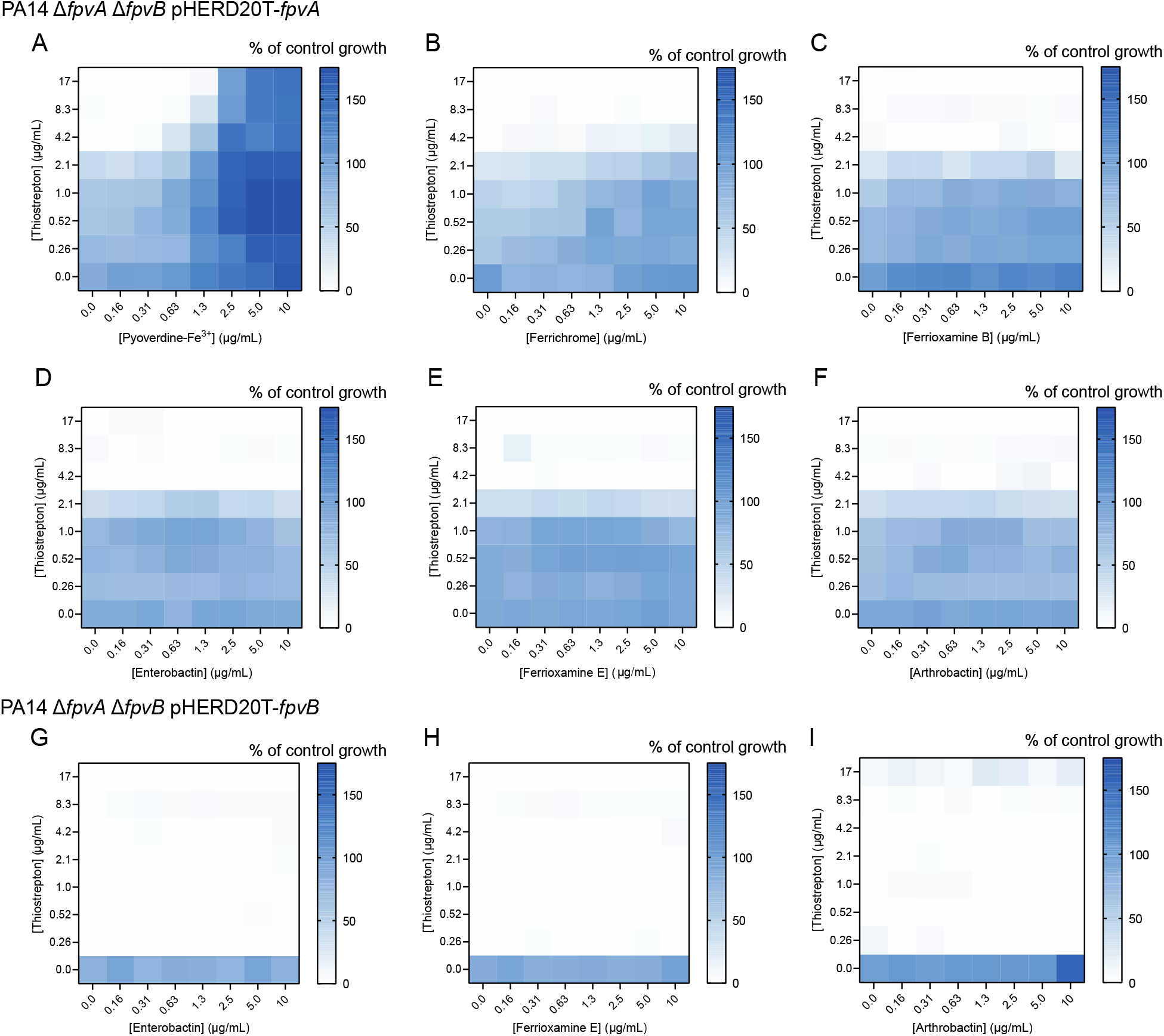
Checkerboard assays of PA14 Δ*fpvA* Δ*fpvB* complemented with *fpvA* and *fpvB in trans* with TS and siderophores. Checkerboards of PA14 Δ*fpvA* Δ*fpvB* pHERD20T-*fpvA* with TS and (A) pyoverdine-Fe^3+^, (B) ferrichrome, (C) ferrioxamine B, (D) enterobactin, (E) ferrioxamine E, and (F) arthrobactin. Checkerboards of PA14 Δ*fpvA* Δ*fpvB* pHERD20T-*fpvB* with TS and (G) enterobactin, (H) ferrioxamine E, and (I) arthrobactin. Results are averaged from three independent biological replicates. Cells in the assay were grown in 10:90 + 2% arabinose.

**Figure 3 – figure supplement 1.**
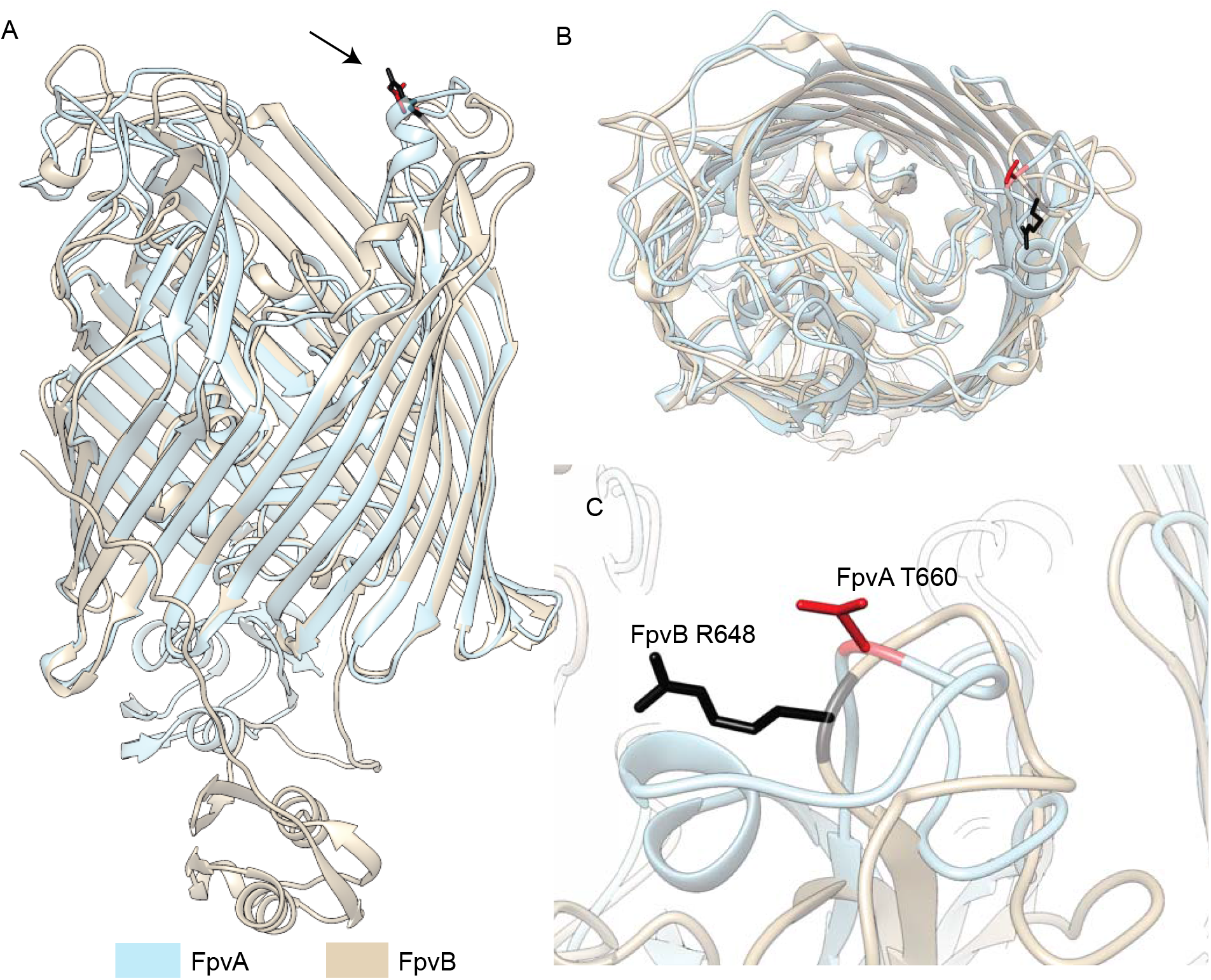
Overlay of FpvA and FpvB showing residues mutated to cysteine for labeling with maleimide dyes. FpvA (PDB: 2O5P) was superimposed with the AlphaFold2 model of FpvB. FpvA is shown in beige and FpvB in blue. (A) Side view, (B) top view, and (C) zoomed in view of the labeled extracellular loop 8.

**Figure 3 – figure supplement 2.**
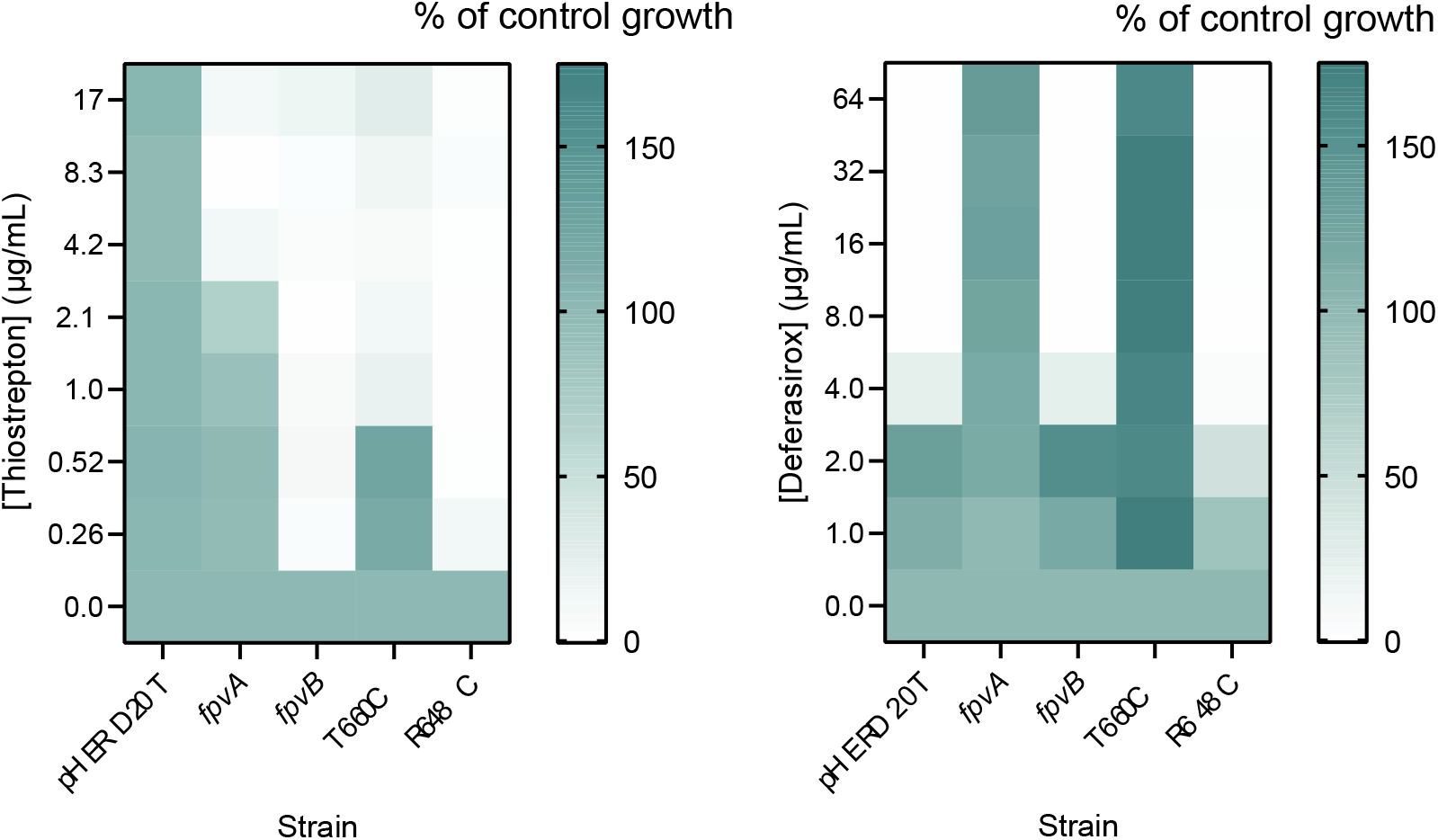
FpvB R648C growth with TS or DSX is similar to WT. FpvA T660C increases susceptibility to TS by 4-fold and restores growth with DSX. Results are averaged from three independent biological replicates. Assays were conducted in 10:90 +2% arabinose.

**Figure 3 – figure supplement 3.**
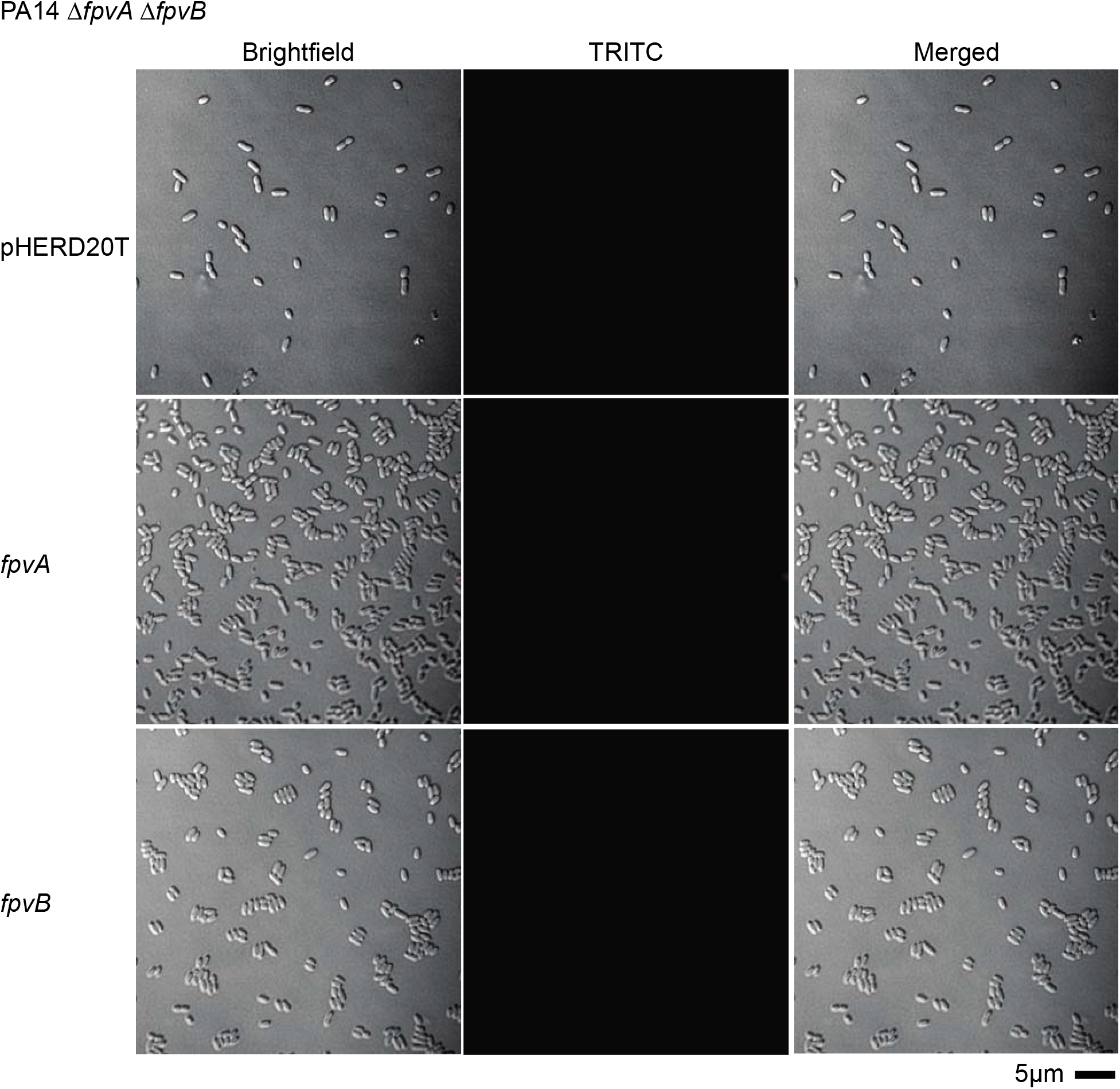
Microscopy images of PA14 Δ*fpvA ΔfpvB* with pHERD20T, pHERD20T-*fpvA*, and pHERD20T-*fpvB* treated with AlexaFluor 594. Representative images are shown. Scale = 5μm.

Ferrichrome-Fe^3+^, ferrioxamine B-Fe^3+^, and pyoverdine-Fe^3+^, were titrated at increasing concentrations into cells expressing FpvA T660C and FpvB R648C labeled with fluorescein-5-maleimide. TS was omitted from these studies because it quenched fluorescence (~20%) of the dye in the absence of cells (Figure 3 – figure supplement 4). In the absence of protein, none of the three siderophores quenched fluorescence of the dye at the highest concentrations tested. Quenching curves were then generated and used to calculate the K_d_. Pyoverdine-Fe^3+^ strongly quenched fluorescence of cells expressing FpvA T660C, with a K_d_ of 10 ± 1.6 ng/mL (8.2 ± 1.2 nM), similar to values reported in previous studies (Figure 3 – figure supplement 5) (Greenwald et al., 2009; Kumar et al., 2022). Ferrichrome-Fe^3+^ weakly quenched the fluorescence of cells expressing FpvA T660C (~20%), suggesting that ferrichrome may interact weakly with FpvA. No antagonism was observed between TS and ferrichrome (Figure 2 – figure supplement 3B), suggesting that the competition is minimal. No quenching was observed with ferrioxamine B-_Fe_^3+^.

**Figure 3 – figure supplement 4.**
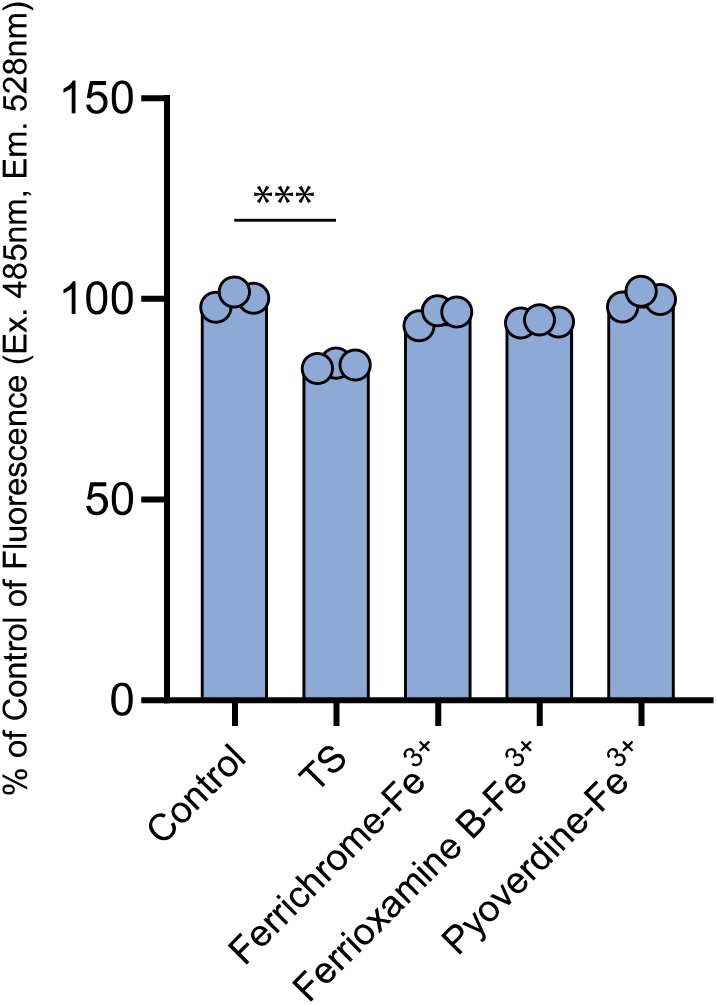
TS partly quenches fluorescence of fluorescein-5-maleimide in the absence of cells. Each compound at the highest concentration tested was added to 10μM of fluorescein-5-maleimide in PBS in triplicate. TS reduced fluorescence to about 80% of the control. Excitation: 485nm; emission 528nm). ***, p<0.001 (student’s t-test).

**Figure 3 – figure supplement 5.**
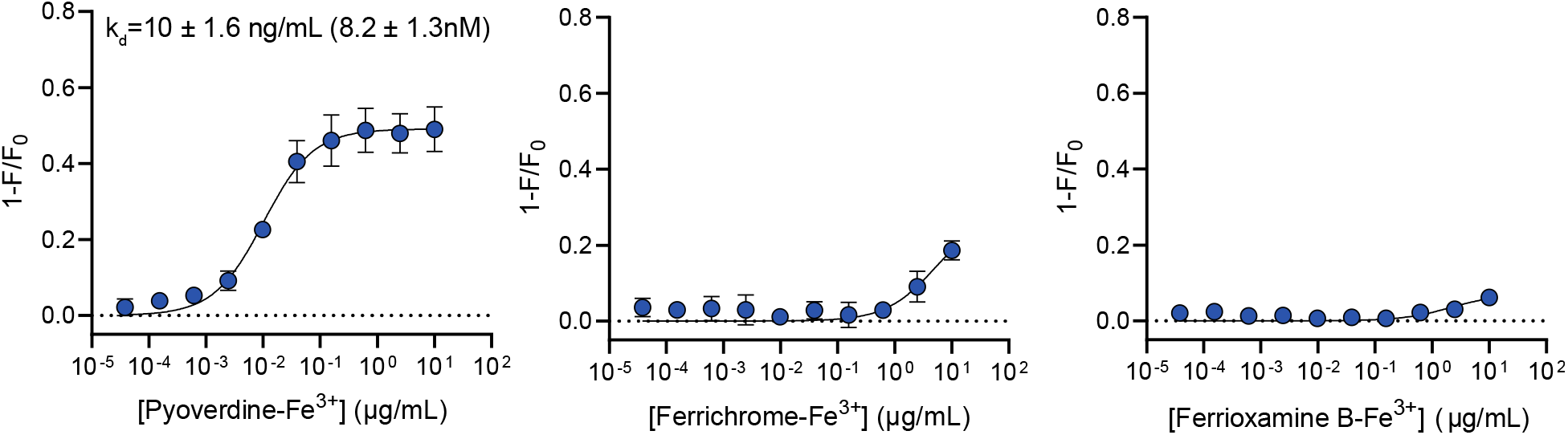
Pyoverdine-Fe^3+^, ferrioxamine B-Fe^3+^ and ferrichrome-Fe^3+^ quenching curves with PA14 Δ*fpvA ΔfpvB* pHERD20T-*fpvA* T660C. Results are averaged from three independent biological replicates.

**Figure 5 – figure supplement 1.**
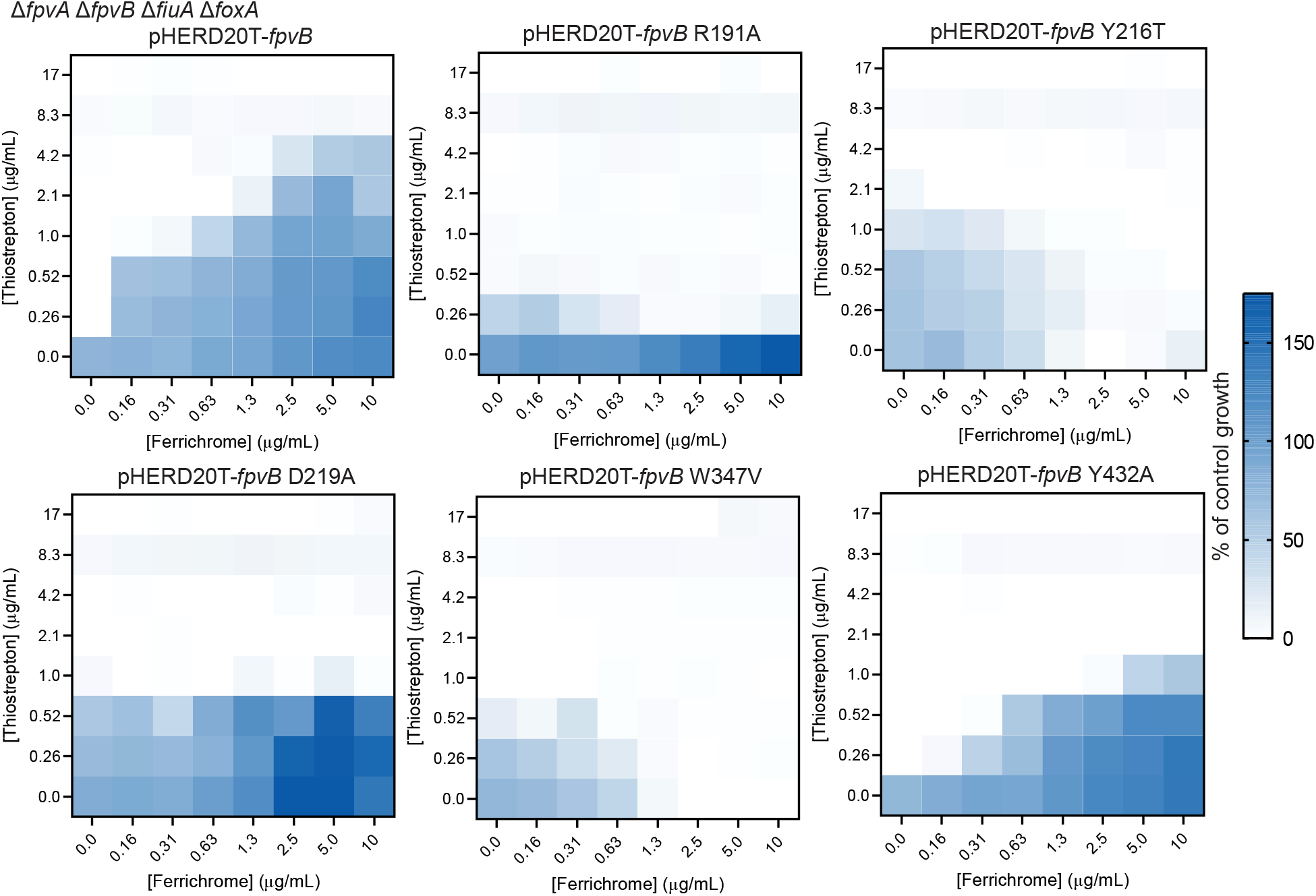
Checkerboard assays in the quadruple transporter mutant background between TS and ferrichrome. Growth is expressed as percent of control. Checkerboards were conducted in 10:90 + 2% arabinose. Results are averaged from three independent biological replicates.

## REFERENCES

Albrecht-Gary AM, Blanc S, Rochel N, Ocaktan AZ, Abdallah MA. 1994. Bacterial Iron Transport: Coordination Properties of Pyoverdin PaA, a Peptidic Siderophore of Pseudomonas aeruginosa. Inorg Chem 33:6391–6402. doi:10.1021/IC00104A059/SUPPL_FILE/IC00104A059_SI_001.PDF

Banin E, Vasil ML, Greenberg EP. 2005. Iron and Pseudomonas aeruginosa biofilm formation. Proc Natl Acad Sci U S A 102:11076–11081. doi:10.1073/PNAS.0504266102/ASSET/4D0E13BE-0F7D-452F-AA17-463C8B67FBA3/ASSETS/GRAPHIC/ZPQ0280588100005.JPEG

Bodilis J, Ghysels B, Osayande J, Matthijs S, Pirnay JP, Denayer S, De Vos D, Cornelis P. 2009. Distribution and evolution of ferripyoverdine receptors in Pseudomonas aeruginosa. Environ Microbiol 11:2123–2135. doi:10.1111/J.1462-2920.2009.01932.X

Braud A, Hannauer M, Mislin GLA, Schalk IJ. 2009. The Pseudomonas aeruginosa pyochelin-iron uptake pathway and its metal specificity. J Bacteriol 191:3517–3525. doi:10.1128/JB.00010-09/ASSET/EEBD944D-0243-472A-A60C-D822F8F24DC8/ASSETS/GRAPHIC/ZJB0110987760005.JPEG

Chakravorty S, Shipelskiy Y, Kumar A, Majumdar A, Yang T, Nairn BL, Newton SM, Klebba PE. 2019. Universal fluorescent sensors of high-affinity iron transport, applied to ESKAPE pathogens. J Biol Chem 294:4682–4692. doi:10.1074/jbc.RA118.006921

Chan DCK, Burrows LL. 2021. Thiocillin and micrococcin exploit the ferrioxamine receptor of Pseudomonas aeruginosa for uptake. J Antimicrob Chemother 76:2029–2039. doi:10.1093/JAC/DKAB124

Chan DCK, Guo I, Burrows LL. 2020. Forging new antibiotic combinations under iron-limiting conditions. Antimicrob Agents Chemother 64. doi:10.1128/AAC.01909-19/SUPPL_FILE/AAC.01909-19-S0001.PDF

Cobessi D, Celia H, Folschweiller N, Schalk IJ, Abdallah MA, Pattus F. 2005. The crystal structure of the pyoverdine outer membrane receptor FpvA from Pseudomonas aeruginosa at 3.6 angstroms resolution. J Mol Biol 347:121–134. doi:10.1016/J.JMB.2005.01.021

Cuív PÓ, Clarke P, O’Connell M. 2006. Identification and characterization of an iron-regulated gene, chtA, required for the utilization of the xenosiderophores aerobactin, rhizobactin 1021 and schizokinen by Pseudomonas aeruginosa. Microbiology 152:945–954. doi:10.1099/MIC.0.28552-0/CITE/REFWORKS

Deatherage DE, Barrick JE. 2014. Identification of mutations in laboratory evolved microbes from next-generation sequencing data using breseq. Methods Mol Biol 1151:165. doi:10.1007/978-1-4939-0554-6_12

Dingemans J, Ye L, Hildebrand F, Tontodonati F, Craggs M, Bilocq F, De Vos D, Crabbé A, Van Houdt R, Malfroot A, Cornelis P. 2014. The deletion of TonB-dependent receptor genes is part of the genome reduction process that occurs during adaptation of Pseudomonas aeruginosa to the cystic fibrosis lung. Pathog Dis 71:26–38. doi:10.1111/2049-632X.12170

Ferguson AD, Coulton JW, Diederichs K, Welte W, Braun V, Fiedler H-P. 2000. Crystal structure of the antibiotic albomycin in complex with the outer membrane transporter FhuA. Protein Sci 9:956–963. doi:10.1110/PS.9.5.956

Ferguson AD, Ködding J, Walker G, Bös C, Coulton JW, Diederichs K, Braun V, Welte W. 2001. Active transport of an antibiotic rifamycin derivative by the outer-membrane protein FhuA. Structure 9:707–716. doi:10.1016/S0969-2126(01)00631-1

Ghysels B, Dieu BTM, Beatson SA, Pirnay JP, Ochsner UA, Vasil ML, Cornelis P. 2004. FpvB, an alternative type I ferripyoverdine receptor of Pseudomonas aeruginosa. Microbiology 150:1671–1680. doi:10.1099/MIC.0.27035-0/CITE/REFWORKS

González J Salvador, Manuel Özkaya Ö, Spick Matt, Reid K Catia Costa, Bailey MJ, Rossa CA, Kümmerli R, Jiménez JI. 2021. Loss of a pyoverdine secondary receptor in Pseudomonas aeruginosa results in a fitter strain suitable for population invasion. ISME J 15:1330–1343. doi:10.1038/s41396-020-00853-2

Greenwald J, Nader M, Celia H, Gruffaz C, Geoffroy V, Meyer JM, Schalk IJ, Pattus F. 2009. FpvA bound to non-cognate pyoverdines: molecular basis of siderophore recognition by an iron transporter. Mol Microbiol 72:1246–1259. doi:10.1111/J.1365-2958.2009.06721.X

Grinter R, Lithgow T. 2019. Determination of the molecular basis for coprogen import by Gram-negative bacteria. IUCrJ 6:401–411. doi:10.1107/S2052252519002926/JT5033SUP12.TXT

Hancock REW, Brinkman FSL. 2003. Function of Pseudomonas Porins in Uptake and Efflux. http://dx.doi.org/101146/annurev.micro56012302160310 56:17–38. doi:10.1146/ANNUREV.MICRO.56.012302.160310

Heinrichs DE, Young L, Poole K. 1991. Pyochelin-mediated iron transport in Pseudomonas aeruginosa: involvement of a high-molecular-mass outer membrane protein. Infect Immun 59:3680–3684. doi:10.1128/IAI.59.10.3680-3684.1991

Hoang TT, Karkhoff-Schweizer RR, Kutchma AJ, Schweizer HP. 1998. A broad-host-range Flp-FRT recombination system for site-specific excision of chromosomally-located DNA sequences: application for isolation of unmarked Pseudomonas aeruginosa mutants. Gene 212:77–86. doi:10.1016/S0378-1119(98)00130-9

Josts I, Veith K, Tidow H. 2019. Ternary structure of the outer membrane transporter FoxA with resolved signaling domain provides insights into TonB-mediated siderophore uptake. Elife 8. doi:10.7554/ELIFE.48528

Jumper J, Evans R, Pritzel A, Green T, Figurnov M, Ronneberger O, Tunyasuvunakool K, Bates R, Žídek A, Potapenko A, Bridgland A, Meyer C, Kohl SAA, Ballard AJ, Cowie A, Romera-Paredes B, Nikolov S, Jain R, Adler J, Back T, Petersen S, Reiman D, Clancy E, Zielinski M, Steinegger M, Pacholska M, Berghammer T, Bodenstein S, Silver D, Vinyals O, Senior AW, Kavukcuoglu K, Kohli P, Hassabis D. 2021. Highly accurate protein structure prediction with AlphaFold. Nat 2021 5967873 596:583–589. doi:10.1038/s41586-021-03819-2

Klebba PE, Newton SMC, Six DA, Kumar A, Yang T, Nairn BL, Munger C, Chakravorty S. 2021. Iron Acquisition Systems of Gram-negative Bacterial Pathogens Define TonB-Dependent Pathways to Novel Antibiotics. Chem Rev 121:5193–5239. doi:10.1021/ACS.CHEMREV.0C01005/SUPPL_FILE/CR0C01005_SI_001.PDF

Koo J, Tammam S, Ku SY, Sampaleanu LM, Burrows LL, Howell PL. 2008. PilF is an outer membrane lipoprotein required for multimerization and localization of the Pseudomonas aeruginosa type IV pilus secretin. J Bacteriol 190:6961–6969. doi:10.1128/JB.00996-08/ASSET/35F08CD4-CEA5-4603-90F7-9401B206E314/ASSETS/GRAPHIC/ZJB0210882100006.JPEG

Kumar A, Yang T, Chakravorty S, Majumdar A, Nairn BL, Six DA, Marcondes N, Santos D, Price SL, Lawrenz MB, Actis LA, Marques M, Russo TA, Newton SM, Klebba PE. 2022. Fluorescent sensors of siderophores produced by bacterial pathogens. J Biol Chem 298:101651. doi:10.1016/j.jbc.2022.101651

Lamont IL, Beare PA, Ochsner U, Vasil AI, Vasil ML. 2002. Siderophore-mediated signaling regulates virulence factor production in Pseudomonas aeruginosa. Proc Natl Acad Sci U S A 99:7072–7077. doi:10.1073/PNAS.092016999/ASSET/65718B21-A04D-4F54-B7F1-89437849CE43/ASSETS/GRAPHIC/PQ0920169005.JPEG

Llamas MA, Sparrius M, Kloet R, Jiménez CR, Vandenbroucke-Grauls C, Bitter W. 2006. The Heterologous Siderophores Ferrioxamine B and Ferrichrome Activate Signaling Pathways in Pseudomonas aeruginosa. J Bacteriol 188:1882. doi:10.1128/JB.188.5.1882-1891.2006

Locher KP, Rees B, Koebnik R, Mitschler A, Moulinier L, Rosenbusch JP, Moras D. 1998. Transmembrane Signaling across the Ligand-Gated FhuA Receptor: Crystal Structures of Free and Ferrichrome-Bound States Reveal Allosteric Changes. Cell 95:771–778. doi:10.1016/S0092-8674(00)81700-6

Luna B, Trebosc V, Lee B, Bakowski M, Ulhaq A, Yan J, Lu P, Cheng J, Nielsen T, Lim J, Ketphan W, Eoh H, McNamara C, Skandalis N, She R, Kemmer C, Lociuro S, Dale GE, Spellberg B. 2020. A nutrient-limited screen unmasks rifabutin hyperactivity for extensively drugresistant Acinetobacter baumannii. Nat Microbiol 2020 59 5:1134–1143. doi:10.1038/s41564-020-0737-6

Luscher A, Moynie L, Auguste P Saint, Bumann D, Mazza L, Pletzer D, Naismith JH, Köhlera T. 2018. TonB-Dependent Receptor Repertoire of Pseudomonas aeruginosa for Uptake of Siderophore-Drug Conjugates. Antimicrob Agents Chemother 62. doi:10.1128/AAC.00097-18

Mathavan I, Zirah S, Mehmood S, Choudhury HG, Goulard C, Li Y, Robinson C V., Rebuffat S, Beis K. 2014. Structural basis for hijacking siderophore receptors by antimicrobial lasso peptides. Nat Chem Biol 2014 105 10:340–342. doi:10.1038/nchembio.1499

Meyer J-M, Geoffroy VA, Baysse C, Cornelis P, Barelmann I, Taraz K, Budzikiewicz H. 2002. Siderophore-mediated iron uptake in fluorescent Pseudomonas: characterization of the pyoverdine-receptor binding site of three cross-reacting pyoverdines. Arch Biochem Biophys 397:179–83. doi:10.1006/abbi.2001.2667

Meyer J-M, Stintzi A, Poole K. 1999. The ferripyoverdine receptor FpvA of Pseudomonas aeruginosa PAO1 recognizes the ferripyoverdines of P. aeruginosa PAO1 and P. fluorescens ATCC 13525. FEMS Microbiol Lett 170:145–150. doi:10.1111/J.1574-6968.1999.TB13367.X

Mirdita M, Schütze K, Moriwaki Y, Heo L, Ovchinnikov S, Steinegger M. 2022. ColabFold: making protein folding accessible to all. Nat Methods 2022 196 19:679–682. doi:10.1038/s41592-022-01488-1

Moynié L, Milenkovic S, Mislin GLA, Gasser V, Malloci G, Baco E, McCaughan RP, Page MGP, Schalk IJ, Ceccarelli M, Naismith JH. 2019. The complex of ferric-enterobactin with its transporter from Pseudomonas aeruginosa suggests a two-site model. Nat Commun 2019 101 10:1–14. doi:10.1038/s41467-019-11508-y

Noinaj N, Guillier M, Barnard TJ, Buchanan SK. 2010. TonB-Dependent Transporters: Regulation, Structure, and Function. http://dx.doi.org/101146/annurev.micro112408134247 64:43–60. doi:10.1146/ANNUREV.MICRO.112408.134247

Normant V, Josts I, Kuhn L, Perraud Q, Fritsch S, Hammann P, Mislin GLA, Tidow H, Schalk IJ. 2020. Nocardamine-Dependent Iron Uptake in Pseudomonas aeruginosa: Exclusive Involvement of the FoxA Outer Membrane Transporter. ACS Chem Biol 15:2741–2751. doi:10.1021/ACSCHEMBIO.0C00535/SUPPL_FILE/CB0C00535_SI_003.XLSX

Ochsner UA, Johnson Z, Vasil ML. 2000. Genetics and regulation of two distinct haem-uptake systems, phu and has, in Pseudomonas aeruginosa. Microbiology 146 (Pt 1):185–198. doi:10.1099/00221287-146-1-185

Parquet M del C, Savage KA, Allan DS, Davidson RJ, Holbein BE. 2018. Novel Iron-Chelator DIBI Inhibits Staphylococcus aureus Growth, Suppresses Experimental MRSA Infection in Mice and Enhances the Activities of Diverse Antibiotics in vitro. Front Microbiol 9. doi:10.3389/FMICB.2018.01811

Pettersen EF, Goddard TD, Huang CC, Couch GS, Greenblatt DM, Meng EC, Ferrin TE. 2004. UCSF Chimera--a visualization system for exploratory research and analysis. J Comput Chem 25:1605–1612. doi:10.1002/JCC.20084

Qiu D, Damron FH, Mima T, Schweizer HP, Yu HD. 2008. PBAD-Based Shuttle Vectors for Functional Analysis of Toxic and Highly Regulated Genes in Pseudomonas and Burkholderia spp. and Other Bacteria. Appl Environ Microbiol 74:7422. doi:10.1128/AEM.01369-08

Rabsch W, Ma L, Wiley G, Najar FZ, Kaserer W, Schuerch DW, Klebba JE, Roe BA, Laverde Gomez JA, Schallmey M, Newton SMC, Klebba PE. 2007. FepA-and TonB-Dependent Bacteriophage H8: Receptor Binding and Genomic Sequence. J Bacteriol 189:5658. doi:10.1128/JB.00437-07

Ranieri MRM, Chan DCK, Yaeger LN, Rudolph M, Karabelas-Pittman S, Abdo H, Chee J, Harvey H, Nguyen U, Burrows LL. 2019. Thiostrepton Hijacks Pyoverdine Receptors to Inhibit Growth of Pseudomonas aeruginosa. Antimicrob Agents Chemother 63. doi:10.1128/AAC.00472-19/SUPPL_FILE/AAC.00472-19-S0003.PDF

Salomon RA, Farias RN. 1993. The FhuA protein is involved in microcin 25 uptake. J Bacteriol 175:7741–7742. doi:10.1128/JB.175.23.7741-7742.1993

Schneider CA, Rasband WS, Eliceiri KW. 2012. NIH Image to ImageJ: 25 years of image analysis. Nat Methods 2012 97 9:671–675. doi:10.1038/nmeth.2089

Shen J, Meldrum A, Poole K. 2002. FpvA Receptor Involvement in Pyoverdine Biosynthesis in Pseudomonas aeruginosa. J Bacteriol 184:3268–3275. doi:10.1128/JB.184.12.3268-3275.2002

Sokol PA. 1987. Tn5 Insertion Mutants of Pseudomonas aeruginosa Deficient in Surface Expression of Ferpyochelin-Binding Protein. J Bacteriol 169:3365–3368.

Southwell JW, Black CM, Duhme-Klair AK. 2021. Experimental Methods for Evaluating the Bacterial Uptake of Trojan Horse Antibacterials. ChemMedChem 16:1063–1076. doi:10.1002/CMDC.202000806

Stefánsson A. 2007. Iron(III) hydrolysis and solubility at 25°C. Environ Sci Technol 41:6117–6123. doi:10.1021/ES070174H

Trott O, Olson AJ. 2010. AutoDock Vina: improving the speed and accuracy of docking with a new scoring function, efficient optimization and multithreading. J Comput Chem 31:455. doi:10.1002/JCC.21334

White P, Joshi A, Rassam P, Housden NG, Kaminska R, Goult JD, Redfield C, McCaughey LC, Walker D, Mohammed S, Kleanthous C. 2017. Exploitation of an iron transporter for bacterial protein antibiotic import. Proc Natl Acad Sci U S A 114:12051–12056. doi:10.1073/PNAS.1713741114/SUPPL_FILE/PNAS.201713741SI.PDF

